# Ecological Patterns of Hymenopteran Pollinators in an Andean Urban Area Derived from Participatory Science

**DOI:** 10.64898/2026.05.21.726748

**Authors:** Daniel Velasco-Cedeño, Nelson Miranda-Moyano, Gabriela F. Moya, Diego F. Cisneros-Heredia

## Abstract

Urbanization presents a significant challenge to pollinator communities worldwide, altering ecological dynamics and species interactions. Understanding the impact of urbanization on pollinators is crucial for developing strategies to mitigate pollinator declines and enhance urban biodiversity. This study investigates hymenopteran pollination ecology in the inter-Andean valley of Quito, Ecuador, using participatory science data. Our dataset, consisting of 2113 hymenopteran records and 556 pollination interactions, reveals complex insect-plant relationships in Quito’s urban environment. We found that alien plant species interacted with more pollinator taxa on average, in contrast to the more specialized interactions involving native plant species. Non-native honeybees (*Apis mellifera*) play a dominant role in these pollination networks, strongly shaping overall network structure. Additionally, both native and alien plants acted as pollination hubs, driving important and diverse interactions. The study documents numerous previously unreported pollination interactions, underscoring the value of participatory science in revealing novel ecological insights. Our findings suggest that Quito’s green spaces function as important refuges for urban pollinators. The use of participatory science records proved invaluable for advancing knowledge of urban pollination ecology, despite its inherent limitations. Encouraging native plant cultivation and public awareness can help mitigate pollinator declines in urban settings. This study adds to growing evidence on urban pollinator ecology and highlights participatory science as a practical tool for addressing ecological challenges in cities.

## Introduction

Insects provide multiple ecosystem services that are indispensable for both human societies and natural ecosystems. Among these, pollination is one of the most critical, sustaining biodiversity, agricultural productivity, and ecosystem stability globally (Potts et al., 2016; Porto et al., 2020). Hymenopterans—bees, wasps, and ants—are particularly relevant as some of the most important pollinating taxa (Majewska & Altizer, 2019). In tropical regions, where approximately 94% of plant species depend on animal pollination, the role of insects becomes even more pronounced (Ollerton et al., 2011). Furthermore, several studies have demonstrated a positive relationship between pollinator richness and agricultural yields, underscoring a direct contribution of diverse pollinator communities to food security and rural livelihoods (Yamamoto et al., 2012; Aizen et al., 2020). Yet these systems are increasingly threatened. Habitat transformation, pesticide use, pathogens, invasive species, and climate change are major drivers of pollinator decline worldwide (Potts et al., 2016), with cascading consequences for wild plant reproduction, crop production, livestock, and human well-being.

Urbanization stands out as one of the most pervasive drivers of ecological change affecting pollinator communities. The continuous growth of cities transforms landscapes into mosaics of fragmented habitats, where native and introduced species coexist under novel ecological pressures (Padayachee et al., 2017; Wang et al., 2022). Cities often represent highly heterogeneous environments, characterized by altered land cover, pollution, and reduced habitat connectivity. While these transformations generally pose challenges for pollinators, urban environments can also provide unexpected opportunities. Public parks, residential gardens, green corridors, and remnant natural areas may supply abundant floral resources and nesting habitats, allowing pollinators to persist within human-dominated ecosystems (Matteson et al., 2008; Baldock et al., 2019; Spotswood et al., 2021). In some contexts, urban habitats may even sustain higher pollinator richness than intensively managed agricultural landscapes (Winfree et al., 2007). However, benefits are context dependent. Urban intensification and densification often erode these benefits, leading to reduced diversity of native pollinators, altered species composition, and an increased dominance of a few generalist taxa (Wenzel et al., 2020; Theodorou et al., 2020).

One emblematic example of this process is the European honeybee (*Apis mellifera*), which thrives in cities worldwide due to generalist foraging, high adaptability, and close association with humans (Breeze et al., 2011). Although honeybees provide key crop pollination services, their overwhelming abundance in urban settings can negatively affect native bees and pollination of wild plants (Paini, 2004; Page & Williams, 2022). Reported impacts include competition for floral resources, pathogen transmission, and alteration of plant–pollinator network structures (Vanbergen et al., 2017). For instance, non-native plants such as *Taraxacum officinale* (common dandelion) or *Trifolium repens* (white clover), which are highly attractive to honeybees, may benefit disproportionately from their presence, while native flora can experience reduced visitation and lower reproductive success (Kandori et al., 2009; Muñoz & Cavieres, 2019). Thus, urban pollination networks frequently display both ecological opportunities and threats, with conservation implications that remain poorly understood in tropical cities.

Despite their importance, research on urban pollination ecology has been heavily biased toward temperate ecosystems and developed countries. In the Neotropics, most studies have focused on small groups of species, using either phytocentric approaches (centered on specific plants) or zoocentric approaches (focused on selected animal taxa), which limits the generality of their conclusions (Gutiérrez-Chacón et al., 2018; Sanguinetti & Singer, 2014; Vizentin-Bugoni et al., 2018). In the tropical Andes, vertebrates—particularly hummingbirds—have received far greater attention than insects (Dellinger et al., 2019; Rivadeneira et al., 2020; Crespo et al., 2021; Sonne et al., 2022). In contrast, insect pollination has been overlooked, even though insects are by far the most diverse and abundant pollinator group. Ecuador, one of the world’s most biodiverse countries, has virtually no published studies that examine multispecies plant-insect pollination networks in urban contexts (Silva et al., 2020; Rueda□Uribe et al., 2025). This gap is acute in the tropical Andes, where research on insect biodiversity and ecology has been largely overlooked, especially in inter-andean dry forests (de la Cadena-Mendoza & Ramón-Cabrera, 2023; Rueda□Uribe et al., 2025). Meanwhile, urbanization in the Andes is advancing rapidly, fragmenting habitats and transforming ecological interactions in unique high-altitude landscapes (Rueda□Uribe et al., 2025). Without baseline data, it is difficult to design effective management strategies that incorporate pollinator conservation into urban development.

Participatory science (also termed as citizen science) has emerged as a powerful approach to address such gaps in knowledge. By mobilizing public involvement, participatory science projects generate large-scale datasets with broad taxonomic and geographic coverage (Figure S1), often at scales unattainable by traditional monitoring (Birkin & Goulson, 2015; Mason & Arathi, 2019; Griffiths□Lee et al., 2020). Platforms such as iNaturalist facilitate this process by combining opportunistic observations, automated taxonomic suggestions, and expert validation, yielding increasingly robust biodiversity datasets (Callaghan et al., 2020, 2022). Several recent studies have highlighted the value of these approaches for urban pollinator ecology, including assessments of hummingbird–plant networks in Mexico (Marín-Gómez et al., 2022) and bee diversity in Brazilian cities (Pereira et al., 2024). Such data are particularly valuable in the Neotropics, where limited funding and logistical challenges constrain systematic field surveys. By leveraging participatory science, researchers can explore ecological questions at scales and levels of detail that would otherwise be prohibitive.

Identifying the most visited flowers is the first step towards supporting and restoring plant– pollinator communities (Crespo et al., 2021). In this study, we use participatory science records from iNaturalist to describe the structure and ecological patterns of hymenopteran pollination communities in Quito, Ecuador, a rapidly growing high-Andean city. Specifically, we aim to (1) describe the structure of hymenopteran pollination networks in an urban Andean environment; (2) identify the plants with the most flower-visiting records; and (3) examine the relative contributions of native versus non-native plant species in shaping pollinator communities. By providing the first comprehensive description of insect pollination networks in the inter-Andean valley of Quito, this work addresses a major gap in urban ecology of the tropical Andes, offers insights into the dual role of cities as both threats and refugees for biodiversity, and informs conservation planning, urban green areas management, and pollinator-friendly gardening.

## Methods

### Study Area

The inter-Andean valley of Quito or *Hoya de Guayllabamba* is an intra-Andean basin located in the northern Andes of Ecuador (Stadel, 1990, Alvarado et al., 2014). It is largely isolated from other intra-Andean basins and from the outer Andean slopes by high mountain chains (Gondard, 1976, Terán, 1976, Stadel, 1990). The inter-Andean valley of Quito encompasses several smaller valleys, high plateaus, and hills, at altitudes between 1800 and 3400 m (Terán, 1976). Climate varies mainly as a function of altitude, and orogenic rainfall is important in vegetation distribution. Lower areas towards the center of the inter-Andean valley are arid (<500 mm/year) and warm (16–22ºC). Higher valleys towards the margins are moist (>1000 mm/year) and cold (6–18ºC). Intermediate precipitation (500–1000 mm/year) and temperature (14-16ºC) occur on the eastern hills and along the gorges and ravines.

Anthropogenic impacts have extensively modified all ecosystems of the inter-Andean valley of Quito. Humans have inhabited the region for more than 10 000 years (Nami & Stanford, 2016). Nowadays, the city of Quito has a consolidated urban center of ca. 202,5 km^2^ and an extensive periurban network of ca. 298,7 km^2^ (STHV-MDMQ, 2010, Carrión & Erazo Espinosa, 2012). Native vegetation is reduced in the urban-periurban matrix; restricted to urban gardens dispersed in the city, *Eucalyptus* forests and regenerating shrublands in gorges and gullies, and lawns in urban parks. The few remnants of natural vegetation occur in the ravines and protective forests on city flanks, often in precarious conservation status. In rural areas, semi-natural ecosystems and cultivated and regenerating shrublands and forests occur alongside crops, and most remnants of native ecosystems are degraded (Cisneros-Heredia et al. 2015).

### Data Collection

We created an iNaturalist traditional project (“Pollination | Quito”) and added records of plant-pollinator interactions from the inter-Andean valley of Quito up to 15 July 2022. We reviewed about 6000 records available at the time. As selection criteria, we included observations with insects in contact with or close to the plant’s sexual organs. We excluded “nectar robbing” events (Maloof & Inouye, 2000). Records belonged both to “Needs ID” and “Research Grade” categories. Unlike Marín-Gómez et al. (2022), who restricted analyses to Research Grade bird records (since birds have a deeper studied taxonomy than insects; Troudet et al., 2017), we retain non-Research Grade insect observations to avoid excluding taxa identified only to genus, tribe or family. We also included observations without the “verifiable” category (e.g., those without a date). iNaturalist’s algorithm provides automated taxonomic suggestions and regional experts curate identifications towards the lowest feasible taxonomic level (Callaghan et al., 2022; Van Horn et al., 2018). In this paper, we focused on the Hymenoptera order and exported records as a CSV file in the iNaturalist default sheet format.

We curated and identified both insect and plant records to the lowest taxonomic level (Ratnieks et al., 2016). Most plants and insects were identified to species; and those not identifiable were treated as morphospecies (Derraik et al., 2010). We used identification guides, catalogs, books and the Tropicos online database for taxonomic information and to classify origin as non-native, native or endemic (Cerón, 2015; Jørgensen & León-Yánez, 1999; Heywood, 1985; Fernández et al., 2015; Padilla et al., 2002; Trópicos, 2022). Species identification was constrained by understudying Andean insect taxonomy or common issues derived from heterogeneous participatory science observation (e.g., the pollinator being in the foreground, inadequate visibility of plant structures, low image resolution or insufficient number of images per record. Plant origin classification was assigned at a national scale; thus “native” denotes indigenous to the Ecuadorian territory, but not necessarily native to the inter-Andean valley of Quito. For example, “Arupo” (*Chionanthus pubescens*) is a native tree to Ecuador but originally distributed in the southern Andes of Ecuador and only brought to Quito as an ornamental plant. With complete taxonomic classifications, we analyze the plant species with the highest occurrence and quantified plant-insect interactions into an interaction matrix. We plotted the ecological network using the bipartite R package (Dormann, 2022).

### Data Analysis

We analyzed plant-insect interactions as ecological networks using *bipartite* R package (Dormann, 2022). The *specieslevel()* function provided network metrics (Dormann, 2022). We focused on five statistical indicators: species degree, species strength, weighted betweenness, weighted closeness and the species specificity index (Marín-Gómez et al 2022; Maruyama et al., 2019). Species degree is the sum of partners per species; species strength indicates the sum of dependencies of each species quantifying a species’ relevance across all its partners (Bascompte et al. 2006); betweenness is a value that describes the centrality of a species in the network by its position on the shortest paths between other nodes (Maruyama et al., 2019); closeness also explains the centrality of a species in the network but by its path lengths to other nodes, and specificity acts as a coefficient of interaction variations (Maruyama et al., 2019). These indicators act as descriptors of each species role in the network. To visualize relationships, we performed principal component analyses (PCAs) and plotted plant-insect bipartite networks to visualize the distribution and combination of interactions among species, highlighting interaction strength (connection lines thickness) and species origin (color), using *bipartite* and *igraph* packages (Dormann, 2022; Csárdi & Nepusz, 2006; Figures 1, 2 and S1).

**Figure. 1.**
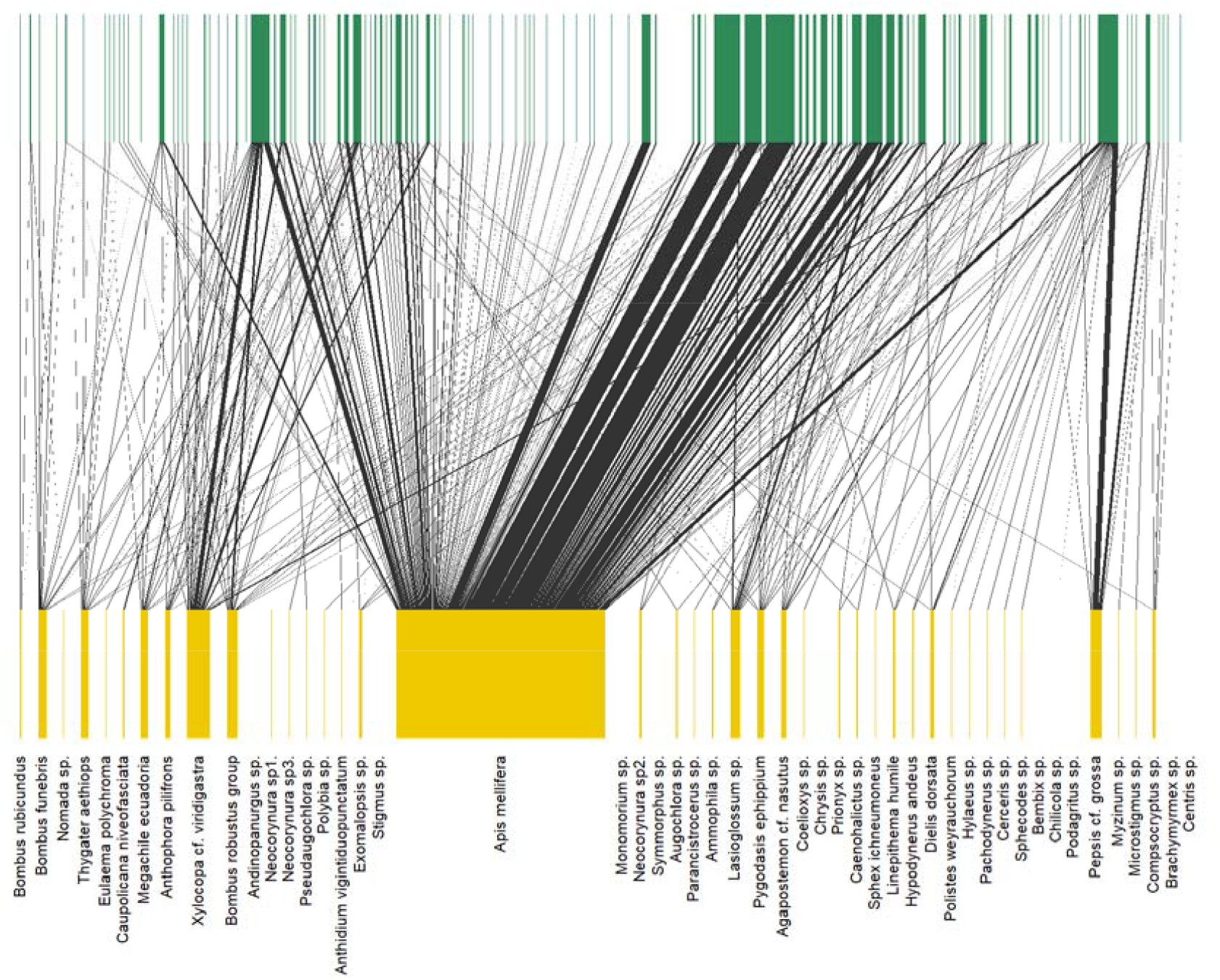
Bipartite Pollination Network. Insect species (yellow blocks) and plant species (dark green blocks) are connected according to pollination interactions. **Alt text:** Bipartite network plot showing interactions between hymenopteran (bottom axis) and plant species (top axis). Each line connects a hymenopteran species to the plant with which it interacts. Yellow blocks at the bottom represent hymenopteran species, with *Apis mellifera* displayed as a wider central bar indicating the highest number of connections. Green blocks at the top represent the associated plants. Black lines illustrate the interactions across the network.

**Figure. 2.**
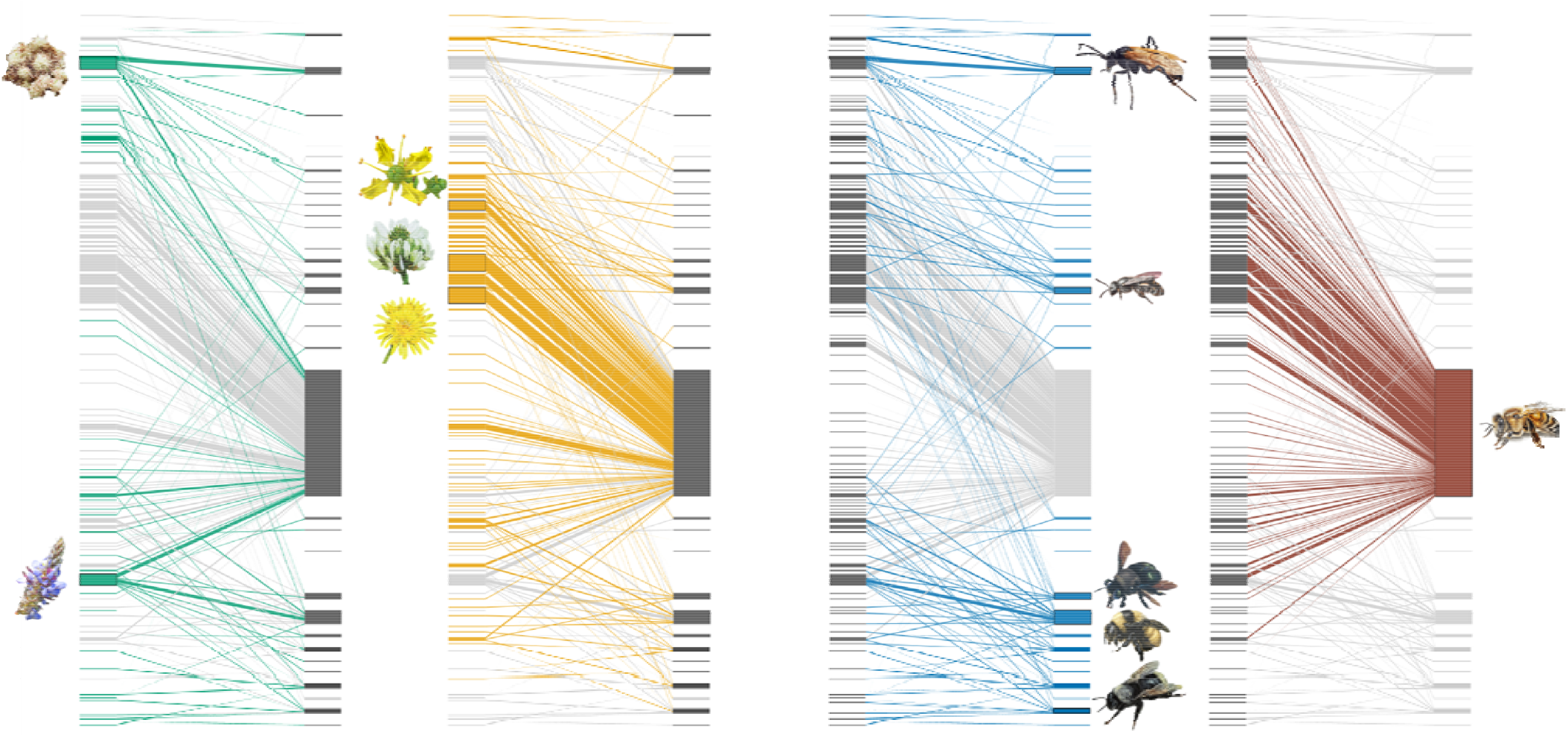
Interactions of Native and Non-native Species. Blue/orange connections denote native/non-native plant interactions; turquoise connections denote native insect interactions; red connections denote *Apis mellifera* interactions. **Alt text:** Bipartite network diagram illustrating interactions between native and non-native plant and insect species. The four panels represent interaction networks with different color codes: turquoise lines indicate native insect interactions, blue lines denote native plant interactions, orange lines denote non-native plant interactions, and red lines represent interactions involving *Apis mellifera*. Each line connects a plant (left side of each panel) with an interacting insect (right side). Images of representative plant and pollinator species are placed alongside their corresponding block.

## Results

Our database comprises 2,113 records (Table S1) and 556 plant-pollinator interactions, including 16 insect families, 46 genera, and 50 species (Table S2), as well as 58 plant families, 147 genera, and 218 species (Table S3). To our knowledge, most of these interactions are unreported in the literature. Non-native taxa accounted for more than half of all plant species and pollination records (128 spp., 66.54% of plant records), while 88 native species (27.78%) were also documented. Ten non-native and three native plant species accounted for 51.63% of all records and were the most frequently visited (Table 1). The most visited plant families were Asteraceae (29.44%), Fabaceae (17.75%), Rutaceae (11.07%), Lamiaceae (9.23%), and Brassicaceae (3.36%). The five most frequently recorded plant species were *Taraxacum officinale* (Asteraceae), *Trifolium repens* (Fabaceae), *Baccharis latifolia* (Asteraceae), *Ruta graveolens* (Rutaceae), and *Dalea coerulea* (Fabaceae) (Figure 1). Among them, *T. officinale, T. repens*, and *R. graveolens* are non-native plants, while *B. latifolia* (locally known as “chilca”) and *D. coerulea* (“iso”) are the most frequently visited native species, acting as key hubs for native pollinators (Figure 1). Less abundant non-native species were *Trifolium pratense* and *Cirsium vulgare* (Table S3).

**Table 1.** Top species of plants by frequency of interactions; origin status indicated.

| # | Family | Species | Common Name | Origin | Freq. |
| --- | --- | --- | --- | --- | --- |
| 1 | Asteraceae | *Taraxacum officinale* | Common dandelion | Alien | 7.71% |
| 2 | Fabaceae | *Trifolium repens* | White clover | Alien | 7.57% |
| 3 | Asteraceae | *Baccharis latifolia* | Chilca | Native | 5.73% |
| 4 | Rutaceae | *Ruta graveolens* | Common Rue | Alien | 5.58% |
| 5 | Fabaceae | *Dalea coerulea* | Iso | Native | 4.92% |
| 6 | Rutaceae | *Citrus medica* | Citron | Alien | 4.64% |
| 7 | Asteraceae | *Hypochaeris radicata* | False dandelion | Alien | 2.93% |
| 8 | Aizoaceae | *Aptenia cordifolia* | Red aptenia | Alien | 2.51% |
| 9 | Lamiaceae | *Salvia microphylla* | Baby sage | Alien | 2.13% |
| 10 | Asteraceae | *Bidens pilosa* | Black-jack | Native | 2.04% |
| 11 | Lamiaceae | *Leonotis nepetifolia* | Lion's ear | Alien | 2.04% |
| 12 | Asteraceae | *Argyranthemum frutescens* | Paris daisy | Alien | 1.94% |
| 13 | Brassicaceae | *Raphanus sativus* | Radish | Alien | 1.89% |
| 14 | Lamiaceae | *Salvia leucantha* | Mexican bush sage | Alien | 1.80% |
| 15 | Asteraceae | *Bidens andicola* | Ñachag | Native | 1.61% |
| 16 | Fabaceae | *Trifolium pratense* | Red clover | Alien | 1.56% |
| 17 | Asteraceae | *Dahlia pinnata* | Garden dahlia | Alien | 1.51% |
| 18 | Apiaceae | *Foeniculum vulgare* | Fennel | Alien | 1.42% |
| 19 | Rosaceae | *Rubus niveus* | Mysore raspberry | Alien | 1.23% |
| 20 | Lythraceae | *Cuphea hyssopifolia* | False heather | Alien | 1.18% |

Insects identified to species or morphospecies accounted for 1,501 records (71.04%); to genus, 1,969 (93.19%); to tribe, 2,047 (96.88%); and to family, 2,100 (99.38%). Plant identifications reached 1,920 records (90.87%) to species, 1,995 (94.42%) to genus, and 2,064 (97.64%) to family. Non-native plants represented 1,406 (66.54%) records and 128 species (58.72%), while native plants comprised 587 records (27.78%) and 88 species (40.37%). Four endemic species (1.83%) were recorded in only five interactions (0.24%), and two species (0.92%) had uncertain origin.

*Apis mellifera* was the most frequent insect species (n = 1,212; 57.36%), followed by *Xylocopa* spp. (n = 130; 6.15%), *Bombus* spp. (n = 115; 5.44%), *Pepsis* spp. (n = 66; 3.12%), *Lasioglossum* spp. (n = 61; 2.89%), and *Megachile* spp. (n = 45; 2.13%). The five most abundant pollinator families were Apidae (73.50%), Halictidae (10.03%), Scoliidae (3.22%), Pompilidae (3.17%), and Vespidae (2.98%). The project includes 924 data contributors (Hymenoptera records), with a single main observer accounting for 17.50% of records; the remaining contributors collectively provided 82.50% of data (, averaging 2.25 records per user). A total of 293 users acted as volunteer taxonomic identifiers, including insect taxonomy experts and local curators (Callaghan et al., 2022).

The ecological networks showed that *A. mellifera* had the highest number of plant partners, interacting with most flowering species (Figure 1). Native pollinators were recorded visiting native plants, including *B. latifolia* and *D. coerulea* (Figure 2). PCAs displayed clustering of most species, with PC1 and PC2 explaining a high proportion of the variance (Figure 3). *Apis mellifera* was analyzed alongside other pollinators in one PCA (Figure 3A) and excluded in another (Figure 3B). PC1 and PC2 separated the five most frequently recorded plant species in analyses with and without non-native species (Figures 3C & D). All variables contributed to PC1 and PC2 scores.

**Figure. 3.**
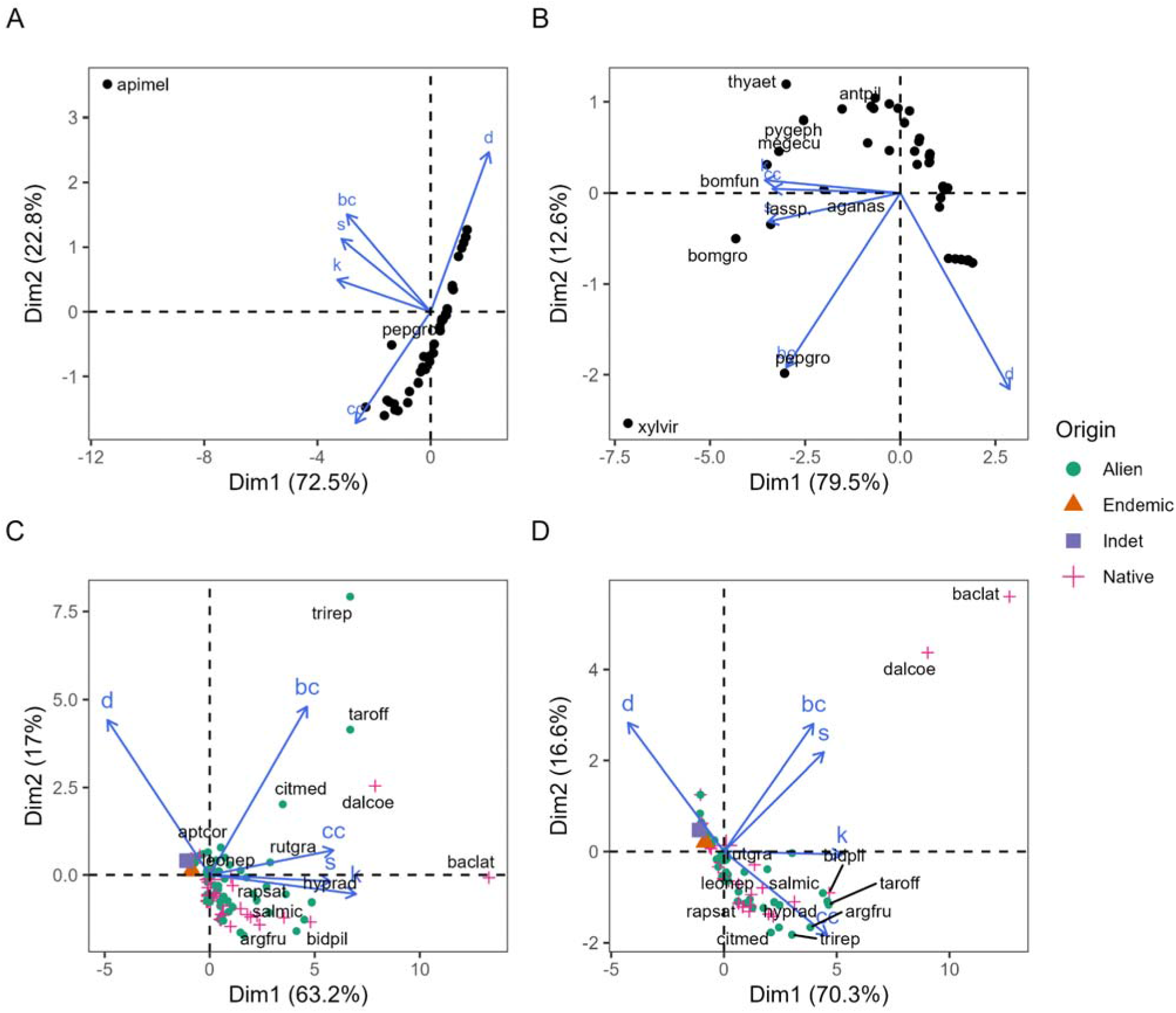
PCA biplots for plant and insect ecological relevance with and without *Apis mellifera* records. Plots A and B show PCAs for insects, including *A. mellifera* in A and excluding them on B, while C and D are PCAs of network indicators for plants, with and without *A. mellifera* in C and D, respectively. Abbreviations correspond to the first three letters of the generic and specific epithets n the respective scientific names. Variables applied to PCAs are network degree (k), strength (s), betweenness centrality (bc), closeness centrality (cc) and species-level specialization (named d’; Dormann, 2022; Maruyama et al., 2019). **Alt text:** Principal Component Analysis (PCA) biplots illustrating the ecological relevance of plant and insect species with and without *Apis mellifera* records included in the analyses. Panels A and B show insect PCAs including (A) and excluding (B) *A. mellifera*, while panels C and D show plant PCAs including (C) and excluding (D) *A. mellifera*. Points represent individual species colored by origin: alien (green), endemic (orange), indeterminate (purple), and native (pink). Blue arrows indicate the direction and contribution of the five network metrics used in the analyses: degree (k), strength (s), betweenness centrality (bc), closeness centrality (cc), and species-level specialization (d’). Each axis label shows the percentage of variance explained by the corresponding principal component. Species are identified by the first three letters of their genus and species epithets.

## Discussion

The western honeybee *Apis mellifera*, a cosmopolitan species introduced into the Neotropics, was the most frequently recorded pollinator in our dataset and the dominant taxon in the bipartite network of Quito. This aligns with studies reporting the remarkable adaptability of honeybees to diverse habitats, including high Andean ecosystems altered by urbanization and agriculture (Knowlton et al., 2022). Honeybees are highly generalist foragers and efficient crop pollinators (Breeze et al., 2011), but their ecological dominance raises important concerns. Their prevalence in the inter-Andean valley is likely linked to anthropogenic land use and managed hives that enhance success in disturbed environments (Ferrer Sánchez et al., 2021). However, interpreting their high record frequency as evidence of ecological primacy is problematic, since observation abundance does not necessarily equate to pollination effectiveness.

Several studies show that honeybees are often less effective pollinators than wild bees on a per-visit basis, with lower pollination efficiency in crops such as cherries and apples (Page & Williams, 2022; Eeraerts et al., 2019). Foraging behaviors, including pollination speed, flower visitation rates, plant-switching probabilities, and stigma contact, frequently differ between honeybees and native pollinators, often favoring the latter (Greenleaf & Kremen, 2006). Consequently, although *A. mellifera* accounted for most recorded interactions in our network, its functional contribution warrants cautious interpretation. In natural and agricultural ecosystems, honeybees may compete for floral resources with native bees, reduce their populations, and indirectly decrease wild plant reproduction (Paini, 2004). Dominance of *A. mellifera* in urban pollination networks of Quito may therefore mask the importance of less abundant but potentially more effective native taxa.

Honeybees interact with a significant proportion of floral species (Figure 1 & 2). Plant species differ in the intensity of honeybee visitation, even within genera. For example, both *Trifolium repens* (white clover) and *Trifolium pratense* (red clover) were recorded as floral resources, but *T. repens* was predominantly visited by honeybees, whereas *T. pratense* was more frequently visited by bumblebees, a pattern also observed in Europe (Kanduth et al., 2021). We also detected *Cirsium vulgare*, a non-native thistle with higher reproductive success than some native congeners (Powell et al., 2010). Among the most abundant species, *Taraxacum officinale* (common dandelion) was heavily visited by both honeybees and native insects (Figure 2), raising concerns about competition with other floral species for pollinator services. Evidence from other regions shows that alien dandelions can reduce reproductive rates of native congeners, such as *T. japonicum* in Japan, by attracting a disproportionate share of pollinators (Kandori et al., 2009). In the Andes, *T. officinale* is a widespread weed competing with native flora for pollination, potentially harming their reproduction and long-term viability in urban environments (Muñoz & Cavieres, 2019). Together, these examples suggest that non-native plants in Quito not only act as key resources for *A. mellifera* but may also disrupt native pollination networks by diverting pollinator visits away from native species.

Our results further highlight strong differentiation between native and non-native plant interactions. Non-native species such as *T. officinale, T. repens*, and *R. graveolens* accounted for a disproportionate share of pollination records. Such plants are highly attractive to generalist pollinators like *A. mellifera*, reinforcing their position as ecological hubs in the network. In contrast, native species such as *B. latifolia* and *D. coerulea* functioned as critical nodes for native bees, underlining their importance in sustaining local pollinator diversity (Watts et al., 2016). The tendency of native insects to preferentially interact with native plants suggests that conserving and promoting native flora in green urban spaces is essential for maintaining functional and resilient pollinator communities in Quito.

The PCAs provided additional insights into species-level relevance. *Apis mellifera* was highly segregated from other pollinators, indicating a distinctive role, while native and non-native plants clustered by their interaction patterns. Importantly, removing honeybee interactions altered the relative position of several species in the PCA space, underscoring the extent to which honeybees structure the network. This pattern is consistent with previous studies showing that invasive pollinators can reshape the topology of plant–pollinator networks, often reducing modularity and increasing asymmetry (Aizen et al., 2008; Vanbergen et al., 2017). Such structural changes may have long-term consequences for ecosystem resilience in fragmented landscapes like the inter-Andean valleys.

Honeybees increase pollination interaction connectivity with high interaction degree, strength, and betweenness centrality (Fig. 1 & 3B), whereas native hymenopterans are more specialised. Native *Baccharis* species are strong hubs, known for their high diversity of flower visitor (Watts et al., 2016). They are visited by several species of bees, wasps and dipterans (Freitas & Sazima, 2006), and, in our data, *B. latifolia* was an important axis visited by spider wasps *Pepsis*, apids, halictids, potter wasps (Eumeninae) and scoliid wasps (Figure 1). *Dalea coerulea* attracted a rich native bee fauna: *Xylocopa* sp., *Bombus* spp., *Anthophora pilifrons, Megachile ecuadoria* and other apoids (Figure 3), being recognised for their appeal to leaf-cutter bees *Megachile* (Figure 1; Cane, 2006). In North América, congeners like *Dalea purpurea* have a similar pollinator composition including honeybees, the alfalfa leafcutting bee (*Megachile rotundata*), and other insects (Cane, 2006).

Beyond species interactions, our results illustrate the dual nature of cities as both threats and refuges for pollinators. On one hand, the prevalence of non-native plants and overwhelming dominance of *A. mellifera* indicate that urbanization facilitates biotic homogenization that can undermine native biodiversity (McKinney, 2006). On the other, the persistence of diverse pollinators, including *Xylocopa, Bombus*, and *Lasioglossum*, demonstrates that urban green spaces retain considerable conservation value. Similar findings in other cities show moderately managed gardens and urban parks can sustain higher pollinator richness than intensively farmed landscapes (Winfree et al., 2007; Baldock et al., 2019). This suggests that the configuration and management of urban green infrastructure are decisive in determining whether cities act as ecological sinks or refugees.

We detected taxonomic and ecological biases typical of participatory science pollination data. Small, elusive, less conspicuous, and less charismatic species were under-recorded (Ratnieks et al., 2016). People usually associate “bees” with the *A. mellifera*, and as a result, more than half (≈57%) of records correspond to that species. This is an obstacle for the monitoring of native bee species, because observers pay more attention to large, highly eusocial bees (Pereira et al., 2024). Nocturnal pollinators are also under-represented, and participatory science observations often recorded them close to light sources or walls and rarely on flowers. The study of the pollination ecology on nocturnal species is almost non-existent in neotropical urban areas, and our study is biased to diurnal records. Nocturnal bees can be efficient crop pollinators, but their systematics and ecology remain poorly studied; only about 1% of described bee species are nocturnal (Cordeiro et al., 2021). Considering light pollution and habitat changes, future urban studies should include nocturnal pollinators (Macgregor & Scott-Brown, 2020).

### Conclusions and recommendations

From a conservation perspective, our findings emphasize integrating pollinator-friendly practices into Quito’s urban planning. Promoting native plants such as *B. latifolia* and *D. coerulea* and other regionally native species in parks, gardens, and restoration projects could enhance resources for native pollinators and mitigate dominance by non-native species. Additionally, awareness campaigns and public engagement are critical, as residents play a direct role in shaping urban floral resources through gardening choices. Previous studies in urban ecology have demonstrated that participatory science not only generates valuable biodiversity data but also fosters environmental stewardship and pro-conservation behaviors among participants (McKinley et al., 2017; Callaghan et al., 2020). Accordingly, our use of iNaturalist records provides ecological insights and exemplifies the potential of participatory monitoring to inform tropical urban management.

Overall, our results underscore the need to promote native flora in green infrastructure and to recognize cities as potential havens for pollinators when appropriately managed.

## Acknowledgements

We thank all iNaturalist users who contributed observations from Quito and made this study possible. We acknowledge use of iNaturalist records under the platform’s Terms of Use and applicable licenses. We thank Jorge Montalvo for taxonomical input on species identification.

## Author contributions

DVC and DFCH conceived and designed the research; DVC collected iNaturalist records with pollination interactions; DVC and DFCH curated insect identification; NMM and GFM curated plant identification; DVC analyzed data and drafted figures; DVC and DFCH drafted the manuscript, and all authors reviewed it.

## Data availability

All data and R code will be made publicly available at https://github.com/davelascoc/pollinators-quito. The database and tables are uploaded at https://doi.org/10.5281/zenodo.20331044.

## Research ethics

This study used publicly available iNaturalist observations; no human subjects were involved, and only non-identifiable metadata were analysed.

## Supporting Information

**Figure S1.**
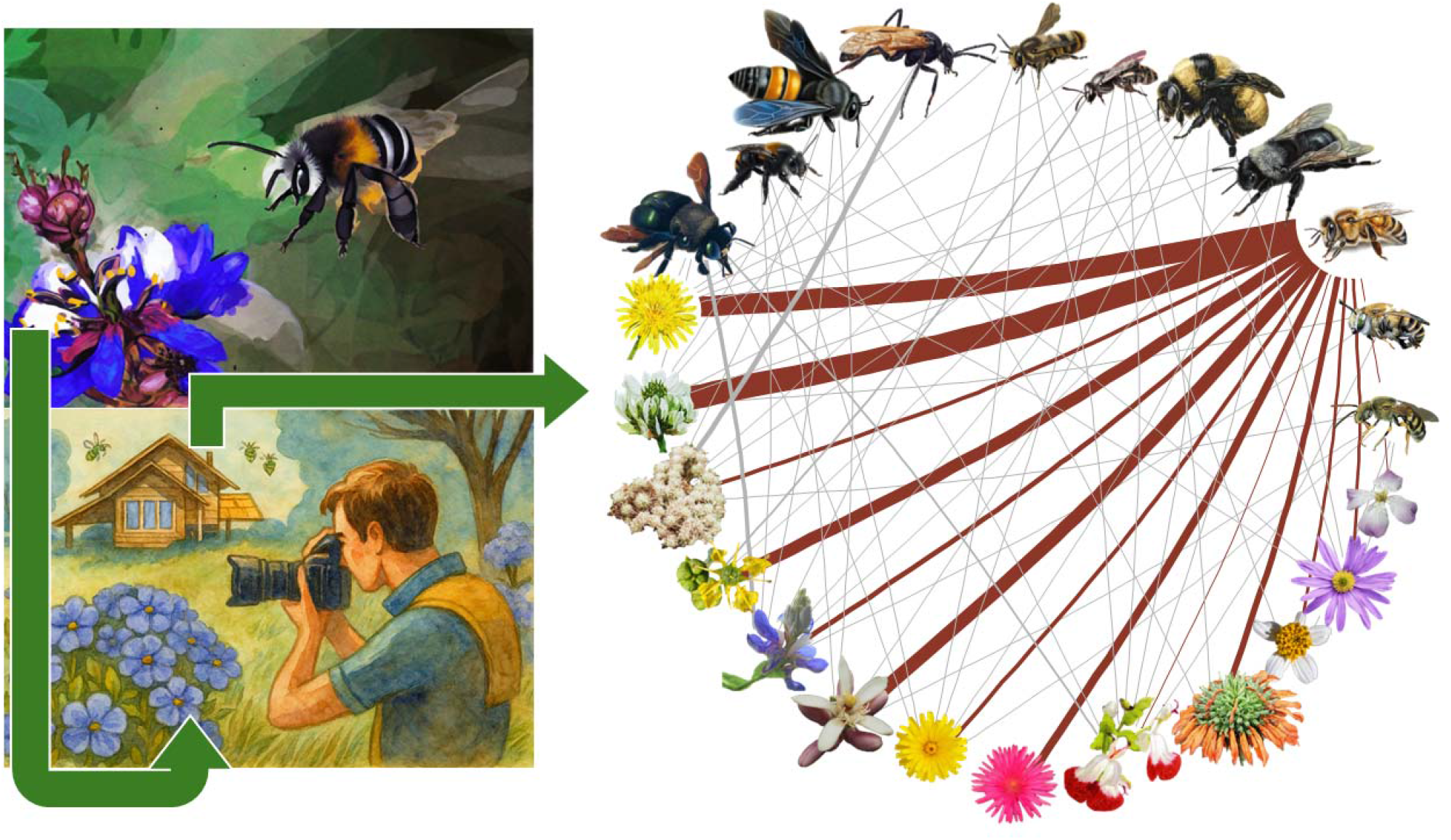
Graphical abstract. **Alt text:** Graphical abstract illustrating the study workflow and main concept. On the left, illustrations depict a native bee visiting a native flowering plant (above), and a person photographing bees on flowers, representing data collection through field observations by participatory scientist (below). On the right, a circular bipartite network shows interactions between hymenopteran species (upper half) and flowering plants (lower half). Red lines represent interaction strength, highlighting *Apis mellifera* as a central and highly connected species within the pollination network. Artistic illustrations were generated using the *Playground* program and the circular ecological network using the *igraph* R package (Csárdi & Nepusz, 2006). Insect pollinators can be observed visiting flowers and citizen scientists generate these records photographing flower-visiting interactions and uploading these records to iNaturalist, where big datasets can be analyzed to describe the pollination community.

