## Supplementary figures and images for "Ecological Patterns of Hymenopteran Pollinators in an Andean Urban Area Derived from Participatory Science"

### Figure S1

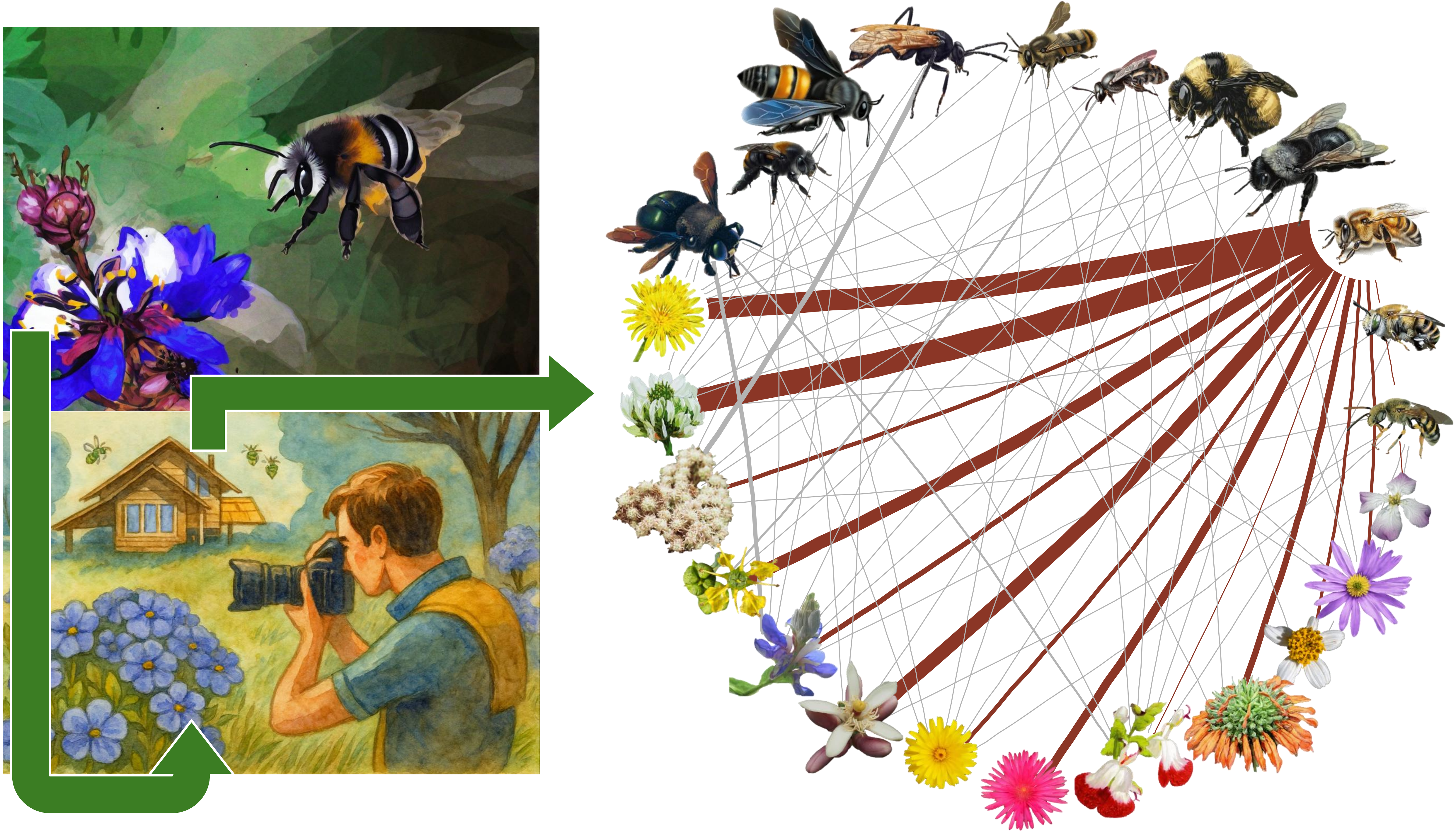
