## Supplementary material for "Ecological Patterns of Hymenopteran Pollinators in an Andean Urban Area Derived from Participatory Science": Table S2

**Table S2.** Complete list of hymenopteran s

| Insect Taxonomy | | | | | | Network Stats | | | | | Network Stats (no Apis) | | | | |
|---|---|---|---|---|---|---|---|---|---|---|---|---|---|---|---|
| **Family** | **Tribe** | **Scientific Name** | **Freq.** | **Rel. Abund.** | **Acr.*** | **k** | **s** | **bc** | **cc** | **d** | **k2** | **s2** | **bc2** | **cc2** | **d2** |
| APIDAE | Apini | *Apis mellifera* | 1212 | 57.36% | apimel | 139 | 103.92 | 0.82 | 0.04 | 0.20 | | | | | |
| Indet | Indet | Indet | 144 | 6.81% | | | | | | | | | | | |
| APIDAE | Xylocopini | *Xylocopa cf. viridigastra* | 130 | 6.15% | xylcf. | 34 | 15.45 | 0.02 | 0.03 | 0.29 | 34 | 21.35 | 0.28 | 0.03 | 0.29 |
| POMPILIDAE | Pepsini | *Pepsis cf. grossa* | 66 | 3.12% | pepcf. | 11 | 4.79 | 0.14 | 0.03 | 0.61 | 11 | 5.99 | 0.19 | 0.03 | 0.61 |
| HALICTIDAE | Halictini | *Lasioglossum sp.* | 61 | 2.89% | lassp. | 20 | 5.03 | 0.00 | 0.03 | 0.34 | 20 | 9.36 | 0.11 | 0.03 | 0.33 |
| APIDAE | Bombini | *Bombus robustus group* | 59 | 2.79% | bomrob | 28 | 7.08 | 0.00 | 0.03 | 0.27 | 28 | 10.95 | 0.14 | 0.03 | 0.26 |
| APIDAE | Bombini | *Bombus funebris* | 48 | 2.27% | bomfun | 23 | 9.99 | 0.02 | 0.02 | 0.26 | 23 | 12.04 | 0.07 | 0.02 | 0.25 |
| MEGACHILIDAE | Megachilini | *Megachile ecuadoria* | 45 | 2.13% | megecu | 22 | 6.83 | 0.00 | 0.02 | 0.29 | 22 | 10.85 | 0.05 | 0.02 | 0.28 |
| APIDAE | Eucerini | *Thygater aethiops* | 42 | 1.99% | thyaet | 24 | 10.26 | 0.00 | 0.02 | 0.24 | 24 | 12.50 | 0.00 | 0.02 | 0.24 |
| SCOLIIDAE | Campsomerini | *Pygodasis ephippium* | 38 | 1.80% | pygeph | 18 | 5.14 | 0.00 | 0.02 | 0.27 | 18 | 9.73 | 0.03 | 0.02 | 0.27 |
| HALICTIDAE | Halictini | *Agapostemon cf. nasutus* | 34 | 1.61% | agacf. | 13 | 2.21 | 0.00 | 0.02 | 0.35 | 13 | 4.86 | 0.08 | 0.02 | 0.34 |
| APIDAE | Anthophorini | *Anthophora pilifrons* | 29 | 1.37% | antpil | 14 | 3.25 | 0.00 | 0.02 | 0.35 | 14 | 4.30 | 0.01 | 0.02 | 0.35 |
| SCOLIIDAE | Campsomerini | *Dielis dorsata* | 19 | 0.90% | | 9 | 1.96 | 0.00 | 0.02 | 0.44 | 9 | 2.46 | 0.02 | 0.02 | 0.44 |
| APIDAE | Exomalopsini | *Exomalopsis sp.* | 17 | 0.80% | | 10 | 1.62 | 0.00 | 0.02 | 0.36 | 10 | 3.12 | 0.01 | 0.02 | 0.36 |
| HALICTIDAE | Augochlorini | *Neocorynura sp2.* | 17 | 0.80% | | 5 | 1.21 | 0.00 | 0.01 | 0.49 | 5 | 1.54 | 0.02 | 0.02 | 0.49 |
| FORMICIDAE | Leptomyrmecini | *Linepithema humile* | 16 | 0.76% | | 10 | 3.13 | 0.00 | 0.01 | 0.33 | 10 | 4.35 | 0.00 | 0.01 | 0.32 |
| ICHNEUMONIDAE | Cryptini | *Compsocryptus sp.* | 16 | 0.76% | | 10 | 4.64 | 0.00 | 0.01 | 0.36 | 10 | 5.23 | 0.00 | 0.01 | 0.36 |
| SPHECIDAE | Ammophilini | *Ammophila sp.* | 10 | 0.47% | | 8 | 1.19 | 0.00 | 0.01 | 0.37 | 8 | 2.49 | 0.00 | 0.01 | 0.37 |
| HALICTIDAE | Halictini | *Caenohalictus sp.* | 9 | 0.43% | | 7 | 0.52 | 0.00 | 0.01 | 0.39 | 7 | 1.74 | 0.00 | 0.01 | 0.39 |
| APIDAE | Bombini | *Bombus rubicundus* | 8 | 0.38% | | 4 | 1.67 | 0.00 | 0.01 | 0.55 | 4 | 2.20 | 0.00 | 0.01 | 0.55 |
| VESPIDAE | Polistini | *Polistes weyrauchorum* | 8 | 0.38% | | 5 | 1.46 | 0.00 | 0.01 | 0.51 | 5 | 1.79 | 0.00 | 0.01 | 0.51 |
| SPHECIDAE | Prionychini | *Prionyx sp.* | 7 | 0.33% | | 6 | 1.95 | 0.00 | 0.01 | 0.42 | 6 | 3.65 | 0.00 | 0.01 | 0.42 |
| VESPIDAE | Eumeninae* | *Hypodynerus andeus* | 7 | 0.33% | | 4 | 0.60 | 0.00 | 0.01 | 0.62 | 4 | 0.67 | 0.00 | 0.01 | 0.62 |
| HALICTIDAE | Augochlorini | *Augochlora sp.* | 6 | 0.28% | | 6 | 0.55 | 0.00 | 0.01 | 0.40 | 6 | 1.00 | 0.00 | 0.01 | 0.40 |
| SPHECIDAE | Sphecini | *Sphex ichneumoneus* | 6 | 0.28% | | 4 | 0.59 | 0.00 | 0.01 | 0.52 | 4 | 0.79 | 0.00 | 0.01 | 0.52 |
| COLLETIDAE | Caupolicanini | *Caupolicana niveofasciata* | 5 | 0.24% | | 1 | 0.05 | 0.00 | 0.01 | 1.00 | 1 | 0.07 | 0.00 | 0.01 | 1.00 |
| COLLETIDAE | Hylaeinae* | *Hylaeus sp.* | 4 | 0.19% | | 2 | 0.10 | 0.00 | 0.01 | 0.79 | 2 | 0.37 | 0.00 | 0.01 | 0.79 |
| HALICTIDAE | Augochlorini | *Pseudaugochlora sp.* | 4 | 0.19% | | 3 | 0.15 | 0.00 | 0.01 | 0.61 | 3 | 0.47 | 0.00 | 0.01 | 0.61 |
| MEGACHILIDAE | Anthidiini | *Anthidium vigintiduopunctatum* | 4 | 0.19% | | 4 | 2.04 | 0.00 | 0.00 | 0.50 | 4 | 2.08 | 0.00 | 0.01 | 0.49 |
| MEGACHILIDAE | Megachilini | *Coelioxys sp.* | 4 | 0.19% | | 3 | 0.07 | 0.00 | 0.01 | 0.61 | 3 | 0.12 | 0.00 | 0.01 | 0.61 |
| APIDAE | Euglossini | *Eulaema polychroma* | 3 | 0.14% | | 3 | 1.17 | 0.00 | 0.00 | 0.57 | 3 | 1.38 | 0.00 | 0.01 | 0.57 |
| APIDAE | Nomadini | *Nomada sp.* | 3 | 0.14% | | 3 | 0.14 | 0.00 | 0.00 | 0.57 | 3 | 0.23 | 0.00 | 0.01 | 0.57 |
| HALICTIDAE | Augochlorini | *Neocorynura sp1.* | 3 | 0.14% | | 3 | 1.03 | 0.00 | 0.00 | 0.57 | 3 | 1.08 | 0.00 | 0.01 | 0.57 |
| HALICTIDAE | Sphecodini | *Sphecodes sp.* | 3 | 0.14% | | 1 | 0.03 | 0.00 | 0.01 | 1.00 | 1 | 0.04 | 0.00 | 0.01 | 1.00 |
| VESPIDAE | Eumeninae* | *Pachodynerus sp.* | 3 | 0.14% | | 2 | 0.52 | 0.00 | 0.00 | 0.74 | 2 | 1.02 | 0.00 | 0.01 | 0.74 |
| APIDAE | Centridini | *Centris sp.* | 2 | 0.09% | | 1 | 1.00 | 0.00 | 0.00 | 1.00 | 1 | 1.00 | 0.00 | 0.00 | 1.00 |
| CRABRONIDAE | Cercerini | *Cerceris sp.* | 2 | 0.09% | | 1 | 0.02 | 0.00 | 0.00 | 1.00 | 1 | 0.02 | 0.00 | 0.01 | 1.00 |
| HALICTIDAE | Augochlorini | *Neocorynura sp3.* | 2 | 0.09% | | 1 | 0.06 | 0.00 | 0.00 | 1.00 | 1 | 0.14 | 0.00 | 0.01 | 1.00 |
| THYNNIDAE | Myzininae* | *Myzinum sp.* | 2 | 0.09% | | 2 | 0.05 | 0.00 | 0.00 | 0.71 | 2 | 0.06 | 0.00 | 0.01 | 0.70 |
| VESPIDAE | Epiponini | *Polybia sp.* | 2 | 0.09% | | 2 | 0.39 | 0.00 | 0.00 | 0.71 | 2 | 1.50 | 0.00 | 0.00 | 0.70 |
| VESPIDAE | Eumeninae* | *Parancistrocerus sp.* | 2 | 0.09% | | 2 | 0.36 | 0.00 | 0.00 | 0.71 | 2 | 0.55 | 0.00 | 0.00 | 0.70 |
| VESPIDAE | Eumeninae* | *Symmorphus sp.* | 2 | 0.09% | | 2 | 0.21 | 0.00 | 0.00 | 0.71 | 2 | 0.39 | 0.00 | 0.01 | 0.70 |
| ANDRENIDAE | Protandrenini | *Andinopanurgus sp.* | 1 | 0.05% | | 1 | 0.25 | 0.00 | 0.00 | 1.00 | 1 | 1.00 | 0.00 | 0.00 | 1.00 |
| CHRYSIDIDAE | Chrysidini | *Chrysis sp.* | 1 | 0.05% | | 1 | 0.25 | 0.00 | 0.00 | 1.00 | 1 | 0.33 | 0.00 | 0.00 | 1.00 |
| COLLETIDAE | Xeromelissinae* | *Chilicola sp.* | 1 | 0.05% | | 1 | 0.01 | 0.00 | 0.00 | 1.00 | 1 | 0.01 | 0.00 | 0.00 | 1.00 |
| CRABRONIDAE | Bembicini | *Bembix sp.* | 1 | 0.05% | | 1 | 0.01 | 0.00 | 0.00 | 1.00 | 1 | 0.01 | 0.00 | 0.00 | 1.00 |

| | | | | | | | | | | | | | | |
|---|---|---|---|---|---|---|---|---|---|---|---|---|---|---|
| CRABRONIDAE | Crabronini | *Podagritus sp.* | 1 | 0.05% | 1 | 0.01 | 0.00 | 0.00 | 1.00 | 1 | 0.01 | 0.00 | 0.00 | 1.00 |
| CRABRONIDAE | Pemphredonini | *Microstigmus sp.* | 1 | 0.05% | 1 | 0.04 | 0.00 | 0.00 | 1.00 | 1 | 0.05 | 0.00 | 0.00 | 1.00 |
| CRABRONIDAE | Pemphredonini | *Stigmus sp.* | 1 | 0.05% | 1 | 0.01 | 0.00 | 0.00 | 1.00 | 1 | 0.06 | 0.00 | 0.00 | 1.00 |
| FORMICIDAE | Myrmelachistini | *Brachymyrmex sp.* | 1 | 0.05% | 1 | 1.00 | 0.00 | 0.00 | 1.00 | 1 | 1.00 | 0.00 | 0.00 | 1.00 |
| FORMICIDAE | Solenopsidini | *Monomorium sp.* | 1 | 0.05% | 1 | 0.03 | 0.00 | 0.00 | 1.00 | 1 | 0.05 | 0.00 | 0.00 | 1.00 |

*Acronyms used in Figure 3 (PCAs)
