## Supplementary material for "Ecological Patterns of Hymenopteran Pollinators in an Andean Urban Area Derived from Participatory Science": Table S3

**Table S3.** Complete list of plant species.

| Plant Taxonomy | | | | | | Network Stats | | | | | Network Stats (no Apis) | | | | |
|---|---|---|---|---|---|---|---|---|---|---|---|---|---|---|---|
| Family | Scientific Name | Origin | Freq. | Rel. Abund. | Acr.* | k | s | bc | cc | d | k2 | s2 | bc2 | cc2 | d2 |
| ASTERACEAE | *Taraxacum officinale* | Alien | 163 | 7.71% | taroff | 13 | 1.63 | 0.17 | 0.01 | 0.84 | 12 | 1.52 | 0.04 | 0.01 | 0.36 |
| FABACEAE | *Trifolium repens* | Alien | 160 | 7.57% | trirep | 8 | 0.57 | 0.30 | 0.01 | 0.89 | 7 | 0.44 | 0.02 | 0.01 | 0.39 |
| ASTERACEAE | *Baccharis latifolia* | Native | 121 | 5.73% | baclat | 23 | 10.08 | 0.10 | 0.01 | 0.39 | 22 | 10.06 | 0.23 | 0.01 | 0.41 |
| Indet | Indet | Indet | 118 | 5.58% | | | | | | | | | | | |
| RUTACEAE | *Ruta graveolens* | Alien | 118 | 5.58% | rutgra | 5 | 1.48 | 0.02 | 0.01 | 0.82 | 4 | 1.41 | 0.01 | 0.01 | 0.83 |
| FABACEAE | *Dalea coerulea* | Native | 104 | 4.92% | dalcoe | 12 | 2.65 | 0.18 | 0.01 | 0.40 | 11 | 2.63 | 0.39 | 0.01 | 0.40 |
| RUTACEAE | *Citrus medica* | Alien | 98 | 4.64% | citmed | 7 | 0.53 | 0.08 | 0.01 | 0.86 | 6 | 0.46 | 0.01 | 0.01 | 0.48 |
| ASTERACEAE | *Hypochaeris radicata* | Alien | 62 | 2.93% | hyprad | 12 | 1.55 | 0.03 | 0.01 | 0.53 | 11 | 1.53 | 0.04 | 0.01 | 0.36 |
| AIZOACEAE | *Aptenia cordifolia* | Alien | 53 | 2.51% | aptcor | 1 | 0.05 | 0.00 | 0.01 | 1.00 | | | | | |
| LAMIACEAE | *Salvia microphylla* | Alien | 45 | 2.13% | salmic | 11 | 1.05 | 0.02 | 0.01 | 0.41 | 10 | 1.04 | 0.06 | 0.01 | 0.46 |
| ASTERACEAE | *Bidens pilosa* | Native | 43 | 2.04% | bidpil | 11 | 2.48 | 0.01 | 0.01 | 0.52 | 10 | 2.47 | 0.03 | 0.01 | 0.33 |
| LAMIACEAE | *Leonotis nepetifolia* | Alien | 43 | 2.04% | leonep | 4 | 0.32 | 0.00 | 0.01 | 0.86 | 3 | 0.28 | 0.00 | 0.01 | 0.61 |
| ASTERACEAE | *Argyranthemum frutescens* | Alien | 41 | 1.94% | argfru | 10 | 1.65 | 0.00 | 0.01 | 0.48 | 9 | 1.63 | 0.00 | 0.01 | 0.36 |
| BRASSICACEAE | *Raphanus sativus* | Alien | 40 | 1.89% | rapsat | 5 | 0.42 | 0.00 | 0.01 | 0.82 | 4 | 0.39 | 0.00 | 0.01 | 0.54 |
| LAMIACEAE | *Salvia leucantha* | Alien | 38 | 1.80% | | 7 | 0.85 | 0.00 | 0.01 | 0.59 | 6 | 0.84 | 0.00 | 0.01 | 0.69 |
| ASTERACEAE | *Bidens andicola* | Native | 34 | 1.61% | | 8 | 1.92 | 0.00 | 0.01 | 0.58 | 7 | 1.90 | 0.00 | 0.01 | 0.37 |
| FABACEAE | *Trifolium pratense* | Alien | 33 | 1.56% | | 5 | 0.41 | 0.04 | 0.01 | 0.52 | 4 | 0.40 | 0.04 | 0.01 | 0.54 |
| ASTERACEAE | *Dahlia pinnata* | Alien | 32 | 1.51% | | 7 | 0.30 | 0.00 | 0.01 | 0.68 | 6 | 0.28 | 0.00 | 0.01 | 0.47 |
| APIACEAE | *Foeniculum vulgare* | Alien | 30 | 1.42% | | 7 | 2.10 | 0.03 | 0.01 | 0.58 | 6 | 2.09 | 0.03 | 0.01 | 0.68 |
| ROSACEAE | *Rubus niveus* | Alien | 26 | 1.23% | | 6 | 0.75 | 0.00 | 0.01 | 0.73 | 5 | 0.73 | 0.00 | 0.01 | 0.50 |
| LYTHRACEAE | *Cuphea hyssopifolia* | Alien | 25 | 1.18% | | 2 | 0.31 | 0.01 | 0.01 | 0.92 | 1 | 0.29 | 0.00 | 0.00 | 1.00 |
| CUCURBITACEAE | *Cucurbita ficifolia* | Alien | 20 | 0.95% | | 3 | 0.55 | 0.01 | 0.00 | 0.89 | 2 | 0.53 | 0.01 | 0.00 | 0.70 |
| FABACEAE | *Phaseolus coccineus* | Native | 19 | 0.90% | | 7 | 0.53 | 0.00 | 0.00 | 0.52 | 6 | 0.53 | 0.00 | 0.01 | 0.59 |
| BRASSICACEAE | *Brassica napus* | Alien | 18 | 0.85% | | 1 | 0.01 | 0.00 | 0.00 | 1.00 | | | | | |
| FABACEAE | *Vicia andicola* | Native | 18 | 0.85% | | 4 | 0.21 | 0.00 | 0.00 | 0.84 | 3 | 0.21 | 0.04 | 0.01 | 0.88 |
| RUTACEAE | *Citrus reticulata* | Alien | 18 | 0.85% | | 2 | 0.04 | 0.00 | 0.01 | 0.94 | 1 | 0.03 | 0.00 | 0.00 | 1.00 |
| MALVACEAE | *Sida poeppigiana* | Native | 17 | 0.80% | | 5 | 0.47 | 0.00 | 0.00 | 0.57 | 4 | 0.47 | 0.00 | 0.01 | 0.69 |
| VERBENACEAE | *Duranta triacantha* | Native | 17 | 0.80% | | 6 | 0.57 | 0.01 | 0.00 | 0.52 | 5 | 0.56 | 0.01 | 0.01 | 0.48 |
| ASTERACEAE | *Helianthus annuus* | Alien | 16 | 0.76% | | 3 | 0.07 | 0.00 | 0.00 | 0.82 | 2 | 0.06 | 0.00 | 0.00 | 0.74 |
| ROSACEAE | *Rubus glaucus* | Native | 16 | 0.76% | | 4 | 0.54 | 0.00 | 0.00 | 0.79 | 3 | 0.53 | 0.00 | 0.00 | 0.57 |
| MYRTACEAE | *Callistemon viminalis* | Alien | 15 | 0.71% | | 2 | 0.26 | 0.00 | 0.00 | 0.87 | 1 | 0.25 | 0.00 | 0.00 | 1.00 |
| OLEACEAE | *Ligustrum japonicum* | Alien | 12 | 0.57% | | 1 | 0.01 | 0.00 | 0.00 | 1.00 | | | | | |
| ASTERACEAE | *Cirsium vulgare* | Alien | 10 | 0.47% | | 3 | 0.22 | 0.00 | 0.00 | 0.67 | 2 | 0.22 | 0.00 | 0.00 | 0.79 |
| ASTERACEAE | *Gazania rigens* | Alien | 10 | 0.47% | | 3 | 0.05 | 0.00 | 0.00 | 0.79 | 2 | 0.04 | 0.00 | 0.00 | 0.70 |
| ASTERACEAE | *Monticalia sp.* | Native | 10 | 0.47% | | 5 | 0.87 | 0.01 | 0.00 | 0.52 | 4 | 0.87 | 0.02 | 0.01 | 0.57 |
| BIGNONIACEAE | *Tecoma stans* | Native | 10 | 0.47% | | 5 | 0.41 | 0.00 | 0.00 | 0.62 | 4 | 0.40 | 0.00 | 0.00 | 0.48 |
| LAMIACEAE | *Lavandula angustifolia* | Alien | 10 | 0.47% | | 3 | 0.08 | 0.00 | 0.00 | 0.70 | 2 | 0.07 | 0.00 | 0.00 | 0.74 |
| VERBENACEAE | *Lantana camara* | Alien | 10 | 0.47% | | 4 | 0.30 | 0.00 | 0.00 | 0.64 | 3 | 0.30 | 0.00 | 0.00 | 0.60 |
| LAURACEAE | *Persea americana* | Native | 9 | 0.43% | | 2 | 0.05 | 0.00 | 0.00 | 0.72 | 1 | 0.05 | 0.00 | 0.00 | 1.00 |
| MYRTACEAE | *Syzygium paniculatum* | Alien | 9 | 0.43% | | 3 | 0.07 | 0.00 | 0.00 | 0.70 | 2 | 0.06 | 0.00 | 0.00 | 0.74 |
| PLANTAGINACEAE | *Hebe speciosa* | Alien | 9 | 0.43% | | 2 | 0.04 | 0.00 | 0.00 | 0.89 | 1 | 0.03 | 0.00 | 0.00 | 1.00 |
| SCROPHULARIACEAE | *Verbascum phlomoides* | Alien | 9 | 0.43% | | 2 | 0.03 | 0.00 | 0.00 | 0.89 | 1 | 0.03 | 0.00 | 0.00 | 1.00 |
| BRASSICACEAE | *Brassica oleracea* | Alien | 8 | 0.38% | | 2 | 0.02 | 0.00 | 0.00 | 0.85 | 1 | 0.02 | 0.00 | 0.00 | 1.00 |
| CRASSULACEAE | *Aeonium canariense* | Alien | 8 | 0.38% | | 1 | 0.01 | 0.00 | 0.00 | 1.00 | | | | | |
| HELIOTROPIACEAE | *Heliotropium arborescens* | Alien | 8 | 0.38% | | 3 | 0.06 | 0.00 | 0.00 | 0.77 | 2 | 0.06 | 0.00 | 0.00 | 0.70 |
| LAMIACEAE | *Rosmarinus officinalis* | Alien | 8 | 0.38% | | 2 | 0.06 | 0.00 | 0.00 | 0.79 | 1 | 0.05 | 0.00 | 0.00 | 1.00 |

| | | | | | | | | | | | | | |
|---|---|---|---|---|---|---|---|---|---|---|---|---|---|
| SOLANACEAE | *Sessea vestita* | Native | 8 | 0.38% | 1 | 0.01 | 0.00 | 0.00 | 1.00 | | | | |
| APIACEAE | *Coriandrum sativum* | Alien | 7 | 0.33% | 4 | 0.07 | 0.00 | 0.00 | 0.61 | 3 | 0.07 | 0.00 | 0.00 | 0.57 |
| ARACEAE | *Zantedeschia aethiopica* | Alien | 7 | 0.33% | 2 | 0.01 | 0.00 | 0.00 | 0.85 | 1 | 0.01 | 0.00 | 0.00 | 1.00 |
| FABACEAE | *Phaseolus vulgaris* | Native | 7 | 0.33% | 3 | 0.06 | 0.00 | 0.00 | 0.65 | 2 | 0.06 | 0.00 | 0.01 | 0.82 |
| AMARYLLIDACEAE | *Allium cepa* | Alien | 6 | 0.28% | 5 | 0.56 | 0.00 | 0.00 | 0.45 | 4 | 0.56 | 0.00 | 0.00 | 0.48 |
| LAMIACEAE | *Minthostachys mollis* | Native | 6 | 0.28% | 2 | 0.09 | 0.00 | 0.00 | 0.85 | 1 | 0.08 | 0.00 | 0.00 | 1.00 |
| PASSIFLORACEAE | *Passiflora ligularis* | Native | 6 | 0.28% | 3 | 0.05 | 0.00 | 0.00 | 0.70 | 2 | 0.05 | 0.00 | 0.00 | 0.70 |
| AMARYLLIDACEAE | *Agapanthus umbellatus* | Alien | 5 | 0.24% | 3 | 0.04 | 0.00 | 0.00 | 0.59 | 2 | 0.04 | 0.00 | 0.00 | 0.74 |
| BORAGINACEAE | *Borago officinalis* | Alien | 5 | 0.24% | 1 | 0.00 | 0.00 | 0.00 | 1.00 | | | | | |
| EUPHORBIACEAE | *Euphorbia pulcherrima* | Alien | 5 | 0.24% | 4 | 1.10 | 0.00 | 0.00 | 0.48 | 3 | 1.10 | 0.00 | 0.00 | 0.57 |
| FABACEAE | *Lupinus pubescens* | Native | 5 | 0.24% | 4 | 0.20 | 0.00 | 0.00 | 0.52 | 4 | 0.20 | 0.00 | 0.01 | 0.51 |
| LAMIACEAE | *Salvia corrugata* | Native | 5 | 0.24% | 2 | 0.05 | 0.00 | 0.00 | 0.82 | 2 | 0.05 | 0.00 | 0.01 | 0.82 |
| LAMIACEAE | *Salvia sp.* | Native | 5 | 0.24% | 3 | 0.12 | 0.00 | 0.00 | 0.59 | 3 | 0.12 | 0.00 | 0.01 | 0.59 |
| LAMIACEAE | *Salvia tiliifolia* | Alien | 5 | 0.24% | 1 | 0.04 | 0.00 | 0.00 | 1.00 | 1 | 0.04 | 0.00 | 0.01 | 1.00 |
| LORANTHACEAE | *Gaiadendron aff. punctatum* | Native | 5 | 0.24% | 2 | 0.08 | 0.00 | 0.00 | 0.71 | 1 | 0.08 | 0.00 | 0.00 | 1.00 |
| MALVACEAE | *Malva dendromorpha* | Alien | 5 | 0.24% | 3 | 0.08 | 0.00 | 0.00 | 0.65 | 2 | 0.08 | 0.00 | 0.01 | 0.79 |
| PLANTAGINACEAE | *Plantago lanceolata* | Alien | 5 | 0.24% | 3 | 0.13 | 0.01 | 0.00 | 0.65 | 2 | 0.12 | 0.00 | 0.00 | 0.70 |
| ROSACEAE | *Hesperomeles sp.* | Native | 5 | 0.24% | 2 | 0.02 | 0.00 | 0.00 | 0.82 | 1 | 0.02 | 0.00 | 0.00 | 1.00 |
| ASPARAGACEAE | *Agave attenuata* | Alien | 4 | 0.19% | 1 | 0.00 | 0.00 | 0.00 | 1.00 | | | | | |
| ASTERACEAE | *Galinsoga parviflora* | Native | 4 | 0.19% | 3 | 0.20 | 0.00 | 0.00 | 0.60 | 2 | 0.20 | 0.00 | 0.00 | 0.70 |
| ASTERACEAE | *Tanacetum parthenium* | Alien | 4 | 0.19% | 3 | 0.52 | 0.00 | 0.00 | 0.57 | 2 | 0.52 | 0.00 | 0.00 | 0.70 |
| BASELLACEAE | *Anredera sp.* | Native | 4 | 0.19% | 2 | 0.06 | 0.00 | 0.00 | 0.79 | 1 | 0.06 | 0.00 | 0.00 | 1.00 |
| BIGNONIACEAE | *Delostoma integrifolium* | Native | 4 | 0.19% | 2 | 0.07 | 0.00 | 0.00 | 0.74 | 2 | 0.07 | 0.00 | 0.00 | 0.74 |
| EUPHORBIACEAE | *Euphorbia milii* | Alien | 4 | 0.19% | 1 | 0.43 | 0.00 | 0.00 | 1.00 | 1 | 0.43 | 0.00 | 0.00 | 1.00 |
| LYTHRACEAE | *Cuphea ignea* | Alien | 4 | 0.19% | 2 | 0.02 | 0.00 | 0.00 | 0.70 | 1 | 0.02 | 0.00 | 0.00 | 1.00 |
| MYRTACEAE | *Melaleuca alternifolia* | Alien | 4 | 0.19% | 1 | 0.00 | 0.00 | 0.00 | 1.00 | | | | | |
| ROSACEAE | *Fragaria vesca* | Alien | 4 | 0.19% | 2 | 1.00 | 0.00 | 0.00 | 0.79 | 1 | 1.00 | 0.00 | 0.00 | 1.00 |
| ROSACEAE | *Rosa centifolia* | Alien | 4 | 0.19% | 1 | 0.00 | 0.00 | 0.00 | 1.00 | | | | | |
| SOLANACEAE | *Lycianthes lycioides* | Native | 4 | 0.19% | 1 | 0.05 | 0.00 | 0.00 | 1.00 | 1 | 0.05 | 0.00 | 0.00 | 1.00 |
| SOLANACEAE | *Nicandra physalodes* | Alien | 4 | 0.19% | 1 | 0.00 | 0.00 | 0.00 | 1.00 | | | | | |
| ASPHODELACEAE | *Aloe vera* | Alien | 3 | 0.14% | 2 | 0.50 | 0.00 | 0.00 | 0.74 | 1 | 0.50 | 0.00 | 0.00 | 1.00 |
| ASPHODELACEAE | *Hemerocallis lilioasphodelus* | Alien | 3 | 0.14% | 1 | 0.17 | 0.00 | 0.00 | 1.00 | 1 | 0.17 | 0.00 | 0.00 | 1.00 |
| ASTERACEAE | *Senecio niveoaureus* | Native | 3 | 0.14% | 1 | 0.07 | 0.00 | 0.00 | 1.00 | 1 | 0.07 | 0.00 | 0.00 | 1.00 |
| BRASSICACEAE | *Raphanus raphanistrum* | Alien | 3 | 0.14% | 1 | 0.00 | 0.00 | 0.00 | 1.00 | | | | | |
| EUPHORBIACEAE | *Euphorbia graminea* | Native | 3 | 0.14% | 2 | 0.33 | 0.00 | 0.00 | 0.70 | 1 | 0.33 | 0.00 | 0.00 | 1.00 |
| FABACEAE | *Inga insignis* | Native | 3 | 0.14% | 1 | 0.00 | 0.00 | 0.00 | 1.00 | | | | | |
| FABACEAE | *Lupinus mutabilis* | Native | 3 | 0.14% | 3 | 0.06 | 0.00 | 0.00 | 0.57 | 3 | 0.06 | 0.00 | 0.00 | 0.57 |
| FABACEAE | *Medicago sativa* | Alien | 3 | 0.14% | 2 | 0.01 | 0.00 | 0.00 | 0.70 | 1 | 0.01 | 0.00 | 0.00 | 1.00 |
| GERANIACEAE | *Pelargonium x hortorum* | Alien | 3 | 0.14% | 2 | 0.05 | 0.00 | 0.00 | 0.70 | 2 | 0.05 | 0.00 | 0.00 | 0.70 |
| GERANIACEAE | *Pelargonium x peltatum* | Alien | 3 | 0.14% | 2 | 0.03 | 0.00 | 0.00 | 0.74 | 1 | 0.03 | 0.00 | 0.00 | 1.00 |
| GERANIACEAE | *Pelargonium zonale* | Alien | 3 | 0.14% | 3 | 0.03 | 0.00 | 0.00 | 0.57 | 2 | 0.02 | 0.00 | 0.00 | 0.70 |
| IRIDACEAE | *Iris florentina* | Alien | 3 | 0.14% | 1 | 0.21 | 0.00 | 0.00 | 1.00 | 1 | 0.21 | 0.00 | 0.00 | 1.00 |
| LAMIACEAE | *Salvia officinalis* | Alien | 3 | 0.14% | 2 | 0.04 | 0.00 | 0.00 | 0.70 | 2 | 0.04 | 0.00 | 0.00 | 0.70 |
| LAMIACEAE | *Salvia sagittata* | Native | 3 | 0.14% | 3 | 0.03 | 0.00 | 0.00 | 0.57 | 2 | 0.03 | 0.00 | 0.00 | 0.70 |
| MELASTOMATACEAE | *Miconia crocea* | Native | 3 | 0.14% | 2 | 0.31 | 0.00 | 0.00 | 0.74 | 2 | 0.31 | 0.00 | 0.00 | 0.74 |
| NYCTAGINACEAE | *Bougainvillea spectabilis* | Alien | 3 | 0.14% | 2 | 0.01 | 0.00 | 0.00 | 0.70 | 1 | 0.01 | 0.00 | 0.00 | 1.00 |
| POACEAE | *Zea mays* | Alien | 3 | 0.14% | 1 | 0.00 | 0.00 | 0.00 | 1.00 | | | | | |
| POLYGALACEAE | *Monnina crassifolia* | Native | 3 | 0.14% | 2 | 0.01 | 0.00 | 0.00 | 0.74 | 1 | 0.01 | 0.00 | 0.00 | 1.00 |
| ROSACEAE | *Eriobotrya japonica* | Alien | 3 | 0.14% | 1 | 0.00 | 0.00 | 0.00 | 1.00 | | | | | |

| | | | | | | | | | | | | | | |
|---|---|---|---|---|---|---|---|---|---|---|---|---|---|---|
| ROSACEAE | *Rosa alba* | Alien | 3 | 0.14% | 2 | 0.01 | 0.00 | 0.00 | 0.74 | 1 | 0.01 | 0.00 | 0.00 | 1.00 |
| SOLANACEAE | *Brugmansia sanguinea* | Native | 3 | 0.14% | 1 | 0.00 | 0.00 | 0.00 | 1.00 | | | | | |
| SOLANACEAE | *Cestrum sp.* | Native | 3 | 0.14% | 2 | 0.14 | 0.00 | 0.00 | 0.74 | 1 | 0.14 | 0.00 | 0.00 | 1.00 |
| SOLANACEAE | *Physalis peruviana* | Native | 3 | 0.14% | 3 | 0.04 | 0.00 | 0.00 | 0.57 | 2 | 0.04 | 0.00 | 0.00 | 0.70 |
| SOLANACEAE | *Solanum furcatum* | Native | 3 | 0.14% | 2 | 0.05 | 0.00 | 0.00 | 0.74 | 1 | 0.05 | 0.00 | 0.00 | 1.00 |
| SOLANACEAE | *Solanum sp.* | Native | 3 | 0.14% | 3 | 0.31 | 0.00 | 0.00 | 0.57 | 2 | 0.31 | 0.00 | 0.00 | 0.70 |
| AMARANTHACEAE | *Alternanthera porrigens* | Native | 2 | 0.09% | 2 | 0.16 | 0.00 | 0.00 | 0.70 | 2 | 0.16 | 0.00 | 0.00 | 0.70 |
| ASTERACEAE | *Ageratina sp.* | Native | 2 | 0.09% | 2 | 0.14 | 0.00 | 0.00 | 0.70 | 1 | 0.14 | 0.00 | 0.00 | 1.00 |
| ASTERACEAE | *Calendula officinalis* | Alien | 2 | 0.09% | 2 | 0.02 | 0.00 | 0.00 | 0.70 | 1 | 0.02 | 0.00 | 0.00 | 1.00 |
| ASTERACEAE | *Conyza bonariensis* | Alien | 2 | 0.09% | 2 | 0.09 | 0.00 | 0.00 | 0.70 | 2 | 0.09 | 0.00 | 0.00 | 0.70 |
| ASTERACEAE | *Gynoxys sp.* | Native | 2 | 0.09% | 2 | 0.10 | 0.00 | 0.00 | 0.70 | 1 | 0.10 | 0.00 | 0.00 | 1.00 |
| ASTERACEAE | *Hypochaeris sessiliflora* | Native | 2 | 0.09% | 1 | 0.04 | 0.00 | 0.00 | 1.00 | 1 | 0.04 | 0.00 | 0.00 | 1.00 |
| ASTERACEAE | *Sonchus oleraceus* | Alien | 2 | 0.09% | 1 | 0.14 | 0.00 | 0.00 | 1.00 | 1 | 0.14 | 0.00 | 0.00 | 1.00 |
| ASTERACEAE | *Tagetes erecta* | Alien | 2 | 0.09% | 2 | 0.02 | 0.00 | 0.00 | 0.70 | 1 | 0.02 | 0.00 | 0.00 | 1.00 |
| ASTERACEAE | *Tagetes sp.* | Alien | 2 | 0.09% | 1 | 0.00 | 0.00 | 0.00 | 1.00 | | | | | |
| BERBERIDACEAE | *Berberis hallii* | Native | 2 | 0.09% | 1 | 0.00 | 0.00 | 0.00 | 1.00 | | | | | |
| CALCEOLARIACEAE | *Calceolaria aff. hyssopifolia* | Native | 2 | 0.09% | 1 | 1.00 | 0.00 | 0.00 | 1.00 | 1 | 1.00 | 0.00 | 0.00 | 1.00 |
| CAPRIFOLIACEAE | *Lonicera japonica* | Alien | 2 | 0.09% | 2 | 0.03 | 0.00 | 0.00 | 0.70 | 1 | 0.03 | 0.00 | 0.00 | 1.00 |
| EUPHORBIACEAE | *Croton elegans* | Endemic | 2 | 0.09% | 2 | 0.10 | 0.00 | 0.00 | 0.70 | 2 | 0.10 | 0.00 | 0.00 | 0.70 |
| FABACEAE | *Medicago sp.* | Alien | 2 | 0.09% | 2 | 0.31 | 0.00 | 0.00 | 0.70 | 2 | 0.31 | 0.00 | 0.00 | 0.70 |
| FABACEAE | *Paraserianthes lophantha* | Alien | 2 | 0.09% | 1 | 0.00 | 0.00 | 0.00 | 1.00 | | | | | |
| GERANIACEAE | *Pelargonium domesticum* | Alien | 2 | 0.09% | 1 | 0.00 | 0.00 | 0.00 | 1.00 | | | | | |
| GERANIACEAE | *Pelargonium x domesticum* | Alien | 2 | 0.09% | 2 | 0.02 | 0.00 | 0.00 | 0.70 | 1 | 0.02 | 0.00 | 0.00 | 1.00 |
| LAMIACEAE | *Salvia aff. hispanica* | Alien | 2 | 0.09% | 2 | 0.06 | 0.00 | 0.00 | 0.70 | 2 | 0.06 | 0.00 | 0.00 | 0.70 |
| LAMIACEAE | *Salvia cf. uliginosa* | Alien | 2 | 0.09% | 2 | 0.03 | 0.00 | 0.00 | 0.70 | 1 | 0.03 | 0.00 | 0.00 | 1.00 |
| LAMIACEAE | *Salvia tortuosa* | Native | 2 | 0.09% | 1 | 0.02 | 0.00 | 0.00 | 1.00 | 1 | 0.02 | 0.00 | 0.00 | 1.00 |
| LAMIACEAE | *Stachys elliptica* | Native | 2 | 0.09% | 2 | 0.01 | 0.00 | 0.00 | 0.70 | 1 | 0.01 | 0.00 | 0.00 | 1.00 |
| MALVACEAE | *Fuertesimalva limensis* | Native | 2 | 0.09% | 2 | 0.12 | 0.00 | 0.00 | 0.70 | 2 | 0.12 | 0.00 | 0.00 | 0.70 |
| NYCTAGINACEAE | *Bougainvillea glabra* | Alien | 2 | 0.09% | 1 | 0.00 | 0.00 | 0.00 | 1.00 | | | | | |
| PAPAVERACEAE | *Papaver sp.* | Alien | 2 | 0.09% | 1 | 0.00 | 0.00 | 0.00 | 1.00 | | | | | |
| POLYGALACEAE | *Monnina phillyreoides* | Alien | 2 | 0.09% | 2 | 0.03 | 0.00 | 0.00 | 0.70 | 2 | 0.03 | 0.00 | 0.00 | 0.70 |
| RUBIACEAE | *Arcytophyllum thymifolium* | Native | 2 | 0.09% | 2 | 0.10 | 0.00 | 0.00 | 0.70 | 1 | 0.10 | 0.00 | 0.00 | 1.00 |
| RUBIACEAE | *Pentas cf. lanceolata* | Alien | 2 | 0.09% | 2 | 0.05 | 0.00 | 0.00 | 0.70 | 2 | 0.05 | 0.00 | 0.00 | 0.70 |
| SOLANACEAE | *Solanum tuberosum* | Native | 2 | 0.09% | 2 | 0.03 | 0.00 | 0.00 | 0.70 | 1 | 0.02 | 0.00 | 0.00 | 1.00 |
| VERBENACEAE | *Aloysia triphylla* | Native | 2 | 0.09% | 1 | 0.00 | 0.00 | 0.00 | 1.00 | | | | | |
| VERBENACEAE | *Verbena litoralis* | Native | 2 | 0.09% | 1 | 0.02 | 0.00 | 0.00 | 1.00 | 1 | 0.02 | 0.00 | 0.00 | 1.00 |
| ACANTHACEAE | *Thunbergia alata* | Alien | 1 | 0.05% | | | | | | | | | | |
| AIZOACEAE | *Lampranthus roseus* | Alien | 1 | 0.05% | 1 | 0.03 | 0.00 | 0.00 | 1.00 | 1 | 0.03 | 0.00 | 0.00 | 1.00 |
| AMARANTHACEAE | *Amaranthus dubius* | Native | 1 | 0.05% | 1 | 0.02 | 0.00 | 0.00 | 1.00 | 1 | 0.02 | 0.00 | 0.00 | 1.00 |
| AMARANTHACEAE | *Chenopodium murale* | Alien | 1 | 0.05% | | | | | | | | | | |
| AMARYLLIDACEAE | *Hippeastrum puniceum* | Native | 1 | 0.05% | | | | | | | | | | |
| AMARYLLIDACEAE | *Ismene longipetala* | Native | 1 | 0.05% | 1 | 0.00 | 0.00 | 0.00 | 1.00 | | | | | |
| ANACARDIACEAE | *Schinus molle* | Alien | 1 | 0.05% | 1 | 0.00 | 0.00 | 0.00 | 1.00 | | | | | |
| APACYNACEAE | *Vinca major* | Alien | 1 | 0.05% | 1 | 0.00 | 0.00 | 0.00 | 1.00 | | | | | |
| ASTERACEAE | *Bellis aff. perennis* | Alien | 1 | 0.05% | 1 | 0.00 | 0.00 | 0.00 | 1.00 | | | | | |
| ASTERACEAE | *Bellis perennis* | Alien | 1 | 0.05% | | | | | | | | | | |
| ASTERACEAE | *Dahlia imperialis* | Alien | 1 | 0.05% | 1 | 0.02 | 0.00 | 0.00 | 1.00 | 1 | 0.02 | 0.00 | 0.00 | 1.00 |
| ASTERACEAE | *Diplostephium ericoides* | Endemic | 1 | 0.05% | 1 | 0.02 | 0.00 | 0.00 | 1.00 | 1 | 0.02 | 0.00 | 0.00 | 1.00 |
| ASTERACEAE | *Gazania sp.* | Alien | 1 | 0.05% | 1 | 0.03 | 0.00 | 0.00 | 1.00 | 1 | 0.03 | 0.00 | 0.00 | 1.00 |

| | | | | | | | | | | | | | | |
|---|---|---|---|---|---|---|---|---|---|---|---|---|---|---|
| ASTERACEAE | *Gnaphalium elegans* | Native | 1 | 0.05% | 1 | 0.33 | 0.00 | 0.00 | 1.00 | 1 | 0.33 | 0.00 | 0.00 | 1.00 |
| ASTERACEAE | *Hypochaeris aff. sessiliflora* | Native | 1 | 0.05% | | | | | | | | | | |
| ASTERACEAE | *Matricaria recutita* | Alien | 1 | 0.05% | | | | | | | | | | |
| ASTERACEAE | *Smallanthus sonchifolius* | Native | 1 | 0.05% | 1 | 0.02 | 0.00 | 0.00 | 1.00 | 1 | 0.02 | 0.00 | 0.00 | 1.00 |
| ASTERACEAE | *Tagetes minuta* | Native | 1 | 0.05% | 1 | 0.14 | 0.00 | 0.00 | 1.00 | 1 | 0.14 | 0.00 | 0.00 | 1.00 |
| ASTERACEAE | *Tagetes patula* | Alien | 1 | 0.05% | 1 | 0.00 | 0.00 | 0.00 | 1.00 | | | | | |
| ASTERACEAE | *Tanacetum sp.* | Alien | 1 | 0.05% | 1 | 0.00 | 0.00 | 0.00 | 1.00 | | | | | |
| ASTERACEAE | *Viguiera quitensis* | Native | 1 | 0.05% | 1 | 0.00 | 0.00 | 0.00 | 1.00 | | | | | |
| BIGNONIACEAE | *Podranea ricasoliana* | Alien | 1 | 0.05% | 1 | 0.02 | 0.00 | 0.00 | 1.00 | 1 | 0.02 | 0.00 | 0.00 | 1.00 |
| BRASSICACEAE | *Capsella bursa-pastoris* | Alien | 1 | 0.05% | 1 | 0.00 | 0.00 | 0.00 | 1.00 | | | | | |
| BRASSICACEAE | *Erucastrum gallicum* | Alien | 1 | 0.05% | | | | | | | | | | |
| CACTACEAE | *Echinopsis pachanoi* | Native | 1 | 0.05% | 1 | 0.00 | 0.00 | 0.00 | 1.00 | | | | | |
| CALCEOLARIACEAE | *Calceolaria crenata* | Native | 1 | 0.05% | 1 | 0.02 | 0.00 | 0.00 | 1.00 | 1 | 0.02 | 0.00 | 0.00 | 1.00 |
| CALCEOLARIACEAE | *Calceolaria sp.* | Native | 1 | 0.05% | 1 | 0.25 | 0.00 | 0.00 | 1.00 | 1 | 0.25 | 0.00 | 0.00 | 1.00 |
| CARYOPHYLLACEAE | *Dianthus caryophyllus* | Alien | 1 | 0.05% | 1 | 0.03 | 0.00 | 0.00 | 1.00 | 1 | 0.03 | 0.00 | 0.00 | 1.00 |
| COMMELINACAEA | *Tradescantia zebrina* | Alien | 1 | 0.05% | | | | | | | | | | |
| CRASSULACEAE | *Kalanchoe densiflora* | Alien | 1 | 0.05% | 1 | 0.06 | 0.00 | 0.00 | 1.00 | 1 | 0.06 | 0.00 | 0.00 | 1.00 |
| CRASSULACEAE | *Sedum sp.* | Alien | 1 | 0.05% | 1 | 0.02 | 0.00 | 0.00 | 1.00 | 1 | 0.02 | 0.00 | 0.00 | 1.00 |
| CUCURBITACEAE | *Cyclanthera pedata* | Native | 1 | 0.05% | 1 | 0.00 | 0.00 | 0.00 | 1.00 | | | | | |
| ESCALLONIACEAE | *Escallonia rubra* | Alien | 1 | 0.05% | 1 | 0.00 | 0.00 | 0.00 | 1.00 | | | | | |
| EUPHORBIACEAE | *Croton aff. pycnanthus* | Endemic | 1 | 0.05% | 1 | 0.02 | 0.00 | 0.00 | 1.00 | 1 | 0.02 | 0.00 | 0.00 | 1.00 |
| EUPHORBIACEAE | *Euphorbia cf. milii* | Alien | 1 | 0.05% | 1 | 1.00 | 0.00 | 0.00 | 1.00 | 1 | 1.00 | 0.00 | 0.00 | 1.00 |
| EUPHORBIACEAE | *Euphorbia leucocephala* | Alien | 1 | 0.05% | 1 | 0.00 | 0.00 | 0.00 | 1.00 | | | | | |
| FABACEAE | *Acasia sp.* | Alien | 1 | 0.05% | 1 | 0.00 | 0.00 | 0.00 | 1.00 | | | | | |
| FABACEAE | *Arachis sp.* | Alien | 1 | 0.05% | 1 | 0.01 | 0.00 | 0.00 | 1.00 | 1 | 0.01 | 0.00 | 0.00 | 1.00 |
| FABACEAE | *Desmodium sp.* | Native | 1 | 0.05% | 1 | 0.07 | 0.00 | 0.00 | 1.00 | 1 | 0.07 | 0.00 | 0.00 | 1.00 |
| FABACEAE | *Lupinus sp.* | Native | 1 | 0.05% | 1 | 0.03 | 0.00 | 0.00 | 1.00 | 1 | 0.03 | 0.00 | 0.00 | 1.00 |
| FABACEAE | *Medicago lupulina* | Alien | 1 | 0.05% | 1 | 0.07 | 0.00 | 0.00 | 1.00 | 1 | 0.07 | 0.00 | 0.00 | 1.00 |
| FABACEAE | *Mimosa albida* | Native | 1 | 0.05% | 1 | 0.00 | 0.00 | 0.00 | 1.00 | | | | | |
| FABACEAE | *Mimosa quitensis* | Native | 1 | 0.05% | 1 | 0.04 | 0.00 | 0.00 | 1.00 | 1 | 0.04 | 0.00 | 0.00 | 1.00 |
| FABACEAE | *Otholobium sp.* | Alien | 1 | 0.05% | 1 | 0.01 | 0.00 | 0.00 | 1.00 | 1 | 0.01 | 0.00 | 0.00 | 1.00 |
| FABACEAE | *Pisum sativum* | Alien | 1 | 0.05% | 1 | 0.02 | 0.00 | 0.00 | 1.00 | 1 | 0.02 | 0.00 | 0.00 | 1.00 |
| FABACEAE | *Senna multiglandulosa* | Native | 1 | 0.05% | | | | | | | | | | |
| FABACEAE | *Senna viarum* | Native | 1 | 0.05% | 1 | 0.02 | 0.00 | 0.00 | 1.00 | 1 | 0.02 | 0.00 | 0.00 | 1.00 |
| FABACEAE | *Vicia faba* | Alien | 1 | 0.05% | 1 | 0.00 | 0.00 | 0.00 | 1.00 | | | | | |
| GENTIANACEAE | *Halenia weddelliana* | Native | 1 | 0.05% | 1 | 0.02 | 0.00 | 0.00 | 1.00 | 1 | 0.02 | 0.00 | 0.00 | 1.00 |
| HYDRANGEACEAE | *Hydrangea macrophylla* | Alien | 1 | 0.05% | 1 | 0.00 | 0.00 | 0.00 | 1.00 | | | | | |
| HYPERICACEAE | *Hypericum perforatum* | Alien | 1 | 0.05% | 1 | 0.00 | 0.00 | 0.00 | 1.00 | | | | | |
| LAMIACEAE | *Prunella vulgaris* | Alien | 1 | 0.05% | 1 | 0.00 | 0.00 | 0.00 | 1.00 | | | | | |
| LAMIACEAE | *Rosmarinus cf. tomentosus* | Alien | 1 | 0.05% | 1 | 0.00 | 0.00 | 0.00 | 1.00 | | | | | |
| LAMIACEAE | *Salvia aff. scutellarioides* | Native | 1 | 0.05% | 1 | 0.00 | 0.00 | 0.00 | 1.00 | | | | | |
| LAMIACEAE | *Salvia cf. hispanica* | Alien | 1 | 0.05% | 1 | 0.01 | 0.00 | 0.00 | 1.00 | 1 | 0.01 | 0.00 | 0.00 | 1.00 |
| LAMIACEAE | *Salvia quitensis* | Endemic | 1 | 0.05% | 1 | 0.02 | 0.00 | 0.00 | 1.00 | 1 | 0.02 | 0.00 | 0.00 | 1.00 |
| MALVACEAE | *Malva sp.* | Indet | 1 | 0.05% | 1 | 0.03 | 0.00 | 0.00 | 1.00 | 1 | 0.03 | 0.00 | 0.00 | 1.00 |
| MYRTACEAE | *Eucalyptus globulus* | Alien | 1 | 0.05% | 1 | 0.02 | 0.00 | 0.00 | 1.00 | 1 | 0.02 | 0.00 | 0.00 | 1.00 |
| ONAGRACEAE | *Fuchsia boliviana* | Alien | 1 | 0.05% | 1 | 0.00 | 0.00 | 0.00 | 1.00 | | | | | |
| ONAGRACEAE | *Fuchsia hybrida* | Alien | 1 | 0.05% | 1 | 0.02 | 0.00 | 0.00 | 1.00 | 1 | 0.02 | 0.00 | 0.00 | 1.00 |
| ONAGRACEAE | *Fuchsia magellanica* | Alien | 1 | 0.05% | 1 | 0.00 | 0.00 | 0.00 | 1.00 | | | | | |
| ONAGRACEAE | *Fuchsia sp.* | Alien | 1 | 0.05% | 1 | 0.33 | 0.00 | 0.00 | 1.00 | 1 | 0.33 | 0.00 | 0.00 | 1.00 |

| | | | | | | | | | | | | | | |
|---|---|---|---|---|---|---|---|---|---|---|---|---|---|---|
| OROBANCHACEAE | *Lamourouxia virgata* | Native | 1 | 0.05% | | | | | | | | | | |
| PASSIFLORACEAE | *Passiflora mixta* | Native | 1 | 0.05% | 1 | 0.00 | 0.00 | 0.00 | 1.00 | | | | | |
| PASSIFLORACEAE | *Passiflora tripartita* | Native | 1 | 0.05% | 1 | 0.00 | 0.00 | 0.00 | 1.00 | | | | | |
| PITTOSPORACEAE | *Pittosporum heterophyllum* | Alien | 1 | 0.05% | 1 | 0.00 | 0.00 | 0.00 | 1.00 | | | | | |
| PITTOSPORACEAE | *Pittosporum undulatum* | Alien | 1 | 0.05% | 1 | 0.00 | 0.00 | 0.00 | 1.00 | | | | | |
| PLANTAGINACEAE | *Veronica persica* | Alien | 1 | 0.05% | | | | | | | | | | |
| POACEAE | *Bromus sp.* | Alien | 1 | 0.05% | 1 | 0.25 | 0.00 | 0.00 | 1.00 | 1 | 0.25 | 0.00 | 0.00 | 1.00 |
| POACEAE | *Eragrostis sp.* | Native | 1 | 0.05% | | | | | | | | | | |
| ROSACEAE | *Malus pumila* | Alien | 1 | 0.05% | 1 | 0.00 | 0.00 | 0.00 | 1.00 | | | | | |
| ROSACEAE | *Prunus serotina* | Native | 1 | 0.05% | 1 | 0.00 | 0.00 | 0.00 | 1.00 | | | | | |
| ROSACEAE | *Rosa sp.* | Alien | 1 | 0.05% | 1 | 0.00 | 0.00 | 0.00 | 1.00 | | | | | |
| ROSACEAE | *Rosa x alba* | Alien | 1 | 0.05% | | | | | | | | | | |
| ROSACEAE | *Rosa x centifolia* | Alien | 1 | 0.05% | 1 | 0.00 | 0.00 | 0.00 | 1.00 | | | | | |
| ROSACEAE | *Rubus adenotrichos* | Native | 1 | 0.05% | 1 | 0.00 | 0.00 | 0.00 | 1.00 | | | | | |
| ROSACEAE | *Rubus bogotensis* | Native | 1 | 0.05% | 1 | 0.00 | 0.00 | 0.00 | 1.00 | | | | | |
| ROSACEAE | *Rubus floribundus* | Native | 1 | 0.05% | 1 | 0.00 | 0.00 | 0.00 | 1.00 | | | | | |
| ROSACEAE | *Rubus idaeus* | Alien | 1 | 0.05% | 1 | 0.00 | 0.00 | 0.00 | 1.00 | | | | | |
| RUBIACEAE | *Psychotria sp.* | Native | 1 | 0.05% | 1 | 0.00 | 0.00 | 0.00 | 1.00 | | | | | |
| SOLANACEAE | *Capsicum rhomboideum* | Native | 1 | 0.05% | 1 | 0.03 | 0.00 | 0.00 | 1.00 | 1 | 0.03 | 0.00 | 0.00 | 1.00 |
| SOLANACEAE | *Datura stramonium* | Native | 1 | 0.05% | 1 | 0.08 | 0.00 | 0.00 | 1.00 | 1 | 0.08 | 0.00 | 0.00 | 1.00 |
| SOLANACEAE | *Solanum lycopersicum* | Native | 1 | 0.05% | 1 | 0.02 | 0.00 | 0.00 | 1.00 | 1 | 0.02 | 0.00 | 0.00 | 1.00 |
| SOLANACEAE | *Solanum quitoense* | Native | 1 | 0.05% | 1 | 0.02 | 0.00 | 0.00 | 1.00 | 1 | 0.02 | 0.00 | 0.00 | 1.00 |
| SOLANACEAE | *Streptosolen jamesonii* | Native | 1 | 0.05% | 1 | 0.01 | 0.00 | 0.00 | 1.00 | 1 | 0.01 | 0.00 | 0.00 | 1.00 |
| TROPAEOLACEAE | *Tropaeolum majus* | Alien | 1 | 0.05% | 1 | 0.00 | 0.00 | 0.00 | 1.00 | | | | | |
| VERBENACEAE | *Lantana achyranthifolia* | Indet | 1 | 0.05% | 1 | 0.05 | 0.00 | 0.00 | 1.00 | 1 | 0.05 | 0.00 | 0.00 | 1.00 |

*Acronyms used in Figure 3 (PCAs)
